# A structural basis for inhibition of the complement initiator protease C1r by Lyme disease spirochetes

**DOI:** 10.1101/2021.01.21.427683

**Authors:** Ryan J. Garrigues, Alexandra D. Powell Pierce, Michal Hammel, Jon T. Skare, Brandon L. Garcia

## Abstract

Complement evasion is a hallmark of extracellular microbial pathogens such as *Borreliella burgdorferi*, the causative agent of Lyme disease. Lyme disease spirochetes express nearly a dozen outer surface lipoproteins that bind complement components and interfere with their native activities. Among these, BBK32 is unique in its selective inhibition of the classical pathway. BBK32 blocks activation of this pathway by selectively binding and inhibiting the C1r serine protease of first component of complement, C1. To understand the structural basis for BBK32-mediated C1r inhibition, we performed co-crystallography and size exclusion chromatography-coupled small angle x-ray scattering experiments, which revealed a molecular model of BBK32-C in complex with activated human C1r. Structure-guided site-directed mutagenesis was combined with surface plasmon resonance binding experiments and assays of complement function to validate the predicted molecular interface. The studies reported here, for the first time, provide a structural basis for classical pathway-specific inhibition by a human pathogen.

## Introduction

The complement system consists of dozens of interacting soluble and surface-associated proteins that together form a primary arm of the innate immune system (*1–4*). Activation of complement on a microbial surface leads to opsonization, recruitment of professional phagocytes, and in many cases, direct cellular lysis. Complement activation involves a cascading series of proteolytic reactions catalyzed by specialized serine proteases that cleave circulating inert complement components into activated fragments. The cascade can be initiated in multiple ways, leading to a canonical grouping of complement activation into three pathways known as the classical, lectin, and alternative pathways. The classical pathway is initiated by a multiprotein complex known as the first component of complement, C1 (*5–8*). C1 is composed of C1q, two molecules of C1r, and two molecules of C1s (C1qr_2_s_2_). C1q serves as the pattern recognition molecule of the classical pathway, binding to immune complexes and non-antibody ligands resulting in zymogen activation of the attached C1r and C1s proteases. The lectin pathway is similarly initiated by pattern recognition, whereby mannan-binding lectin (MBL), ficolins, collectin-10 or collectin-11 bind foreign carbohydrate structures and cause activation of the MBL-associated proteases (MASPs) (*9, 10*). In contrast, the alternative pathway does not rely on pattern recognition but is instead continuously activated at a low level by a process known as ‘tick-over’ (*11*). All three initiating pathways converge at the cleavage of the central molecule of the cascade, complement component C3. C3 activation leads to formation of surface-bound enzyme complexes known as convertases that drive complement amplification and terminal pathway activation, ultimately resulting in the formation of the terminal pathway complete complex, C5b-9, also known as the membrane attack complex (MAC) (*1, 2*).

Complement activation is a finely tuned process controlled by endogenous regulators of complement activity (RCAs) (*12*). RCAs are critical in preventing inappropriate complement activation on the surface of healthy host cells. Indeed, aberrant activation or overactivation of complement causes or exacerbates a wide range of human autoimmune, inflammatory, and neurodegenerative diseases (*13, 14*). Microbes lack RCAs and thus many human pathogens have evolved sophisticated complement evasion systems to prevent complement-mediated attack (*15*). A prototypical example of a bacterial complement evasion system is found in *Borreliella* spirochetes (*16, 17*). *Borreliella* gen. nov. encompasses the etiological agents of Lyme disease including the major genospecies *Borreliella burgdorferi, B. afzelii,* and *B. garinii* (*18*). The *Borreliella* complement evasion repertoire is overlapping and is categorized by two general mechanisms (*16, 17*). The first mechanism involves functional recruitment of host RCAs to the spirochete, surface such as factor H (FH) (*19–28*). Among this group are the complement regulator acquiring surface proteins (CRASPs) which include three distinct classes of FH-binding proteins called CspA (CRASP-1), CspZ (CRASP-2), and three OspEF-related family paralogs known as ErpP (CRASP-3), ErpC (CRASP-4), and ErpA (CRASP-5) (*19–28*). The second general mechanism of complement evasion by outer surface *Borreliella* proteins involves a direct mode of inhibition. Known inhibitors from this group include *B. bavariensis* BGA66 and BGA71 (*29*), a FH-binding independent activity for CspA (*30*), and a protein of unknown molecular identity with CD59-like activity that each interact with components of C5b-9 and prevent MAC formation (*31*). In addition, two *Borreliella* outer surface lipoproteins are known to directly interfere with upstream initiation steps of complement: OspC which binds to C4b and prevents classical/lectin pathway proconvertase formation (*32*), and BBK32 which selectively blocks classical pathway activation (*33*). *B. burgdorferi* BBK32 is an outer surface-localized lipoprotein upregulated during the vertebrate infection phase of the *B. burgdorferi* lifecycle (*34*). We have previously shown that a *bbk32* mutant is significantly attenuated in murine models of infection due to defects in dissemination and the inability of the borrelial cells to maintain a normal bacterial load (*35, 36*). BBK32 contains non-overlapping N-terminal binding sites for host glycosaminoglycans (GAGs) and fibronectin and the contribution of BBK32 to Lyme disease pathogenesis is likely related to its multifunctionality (*37, 38*). Interactions by BBK32 with GAGs and fibronectin at the host endothelium exhibit dragging and tethering, respectively, thereby promoting adherence during to the shear force of blood flow, resulting in the hematogenous dissemination of *B. burgdorferi* via extravasation (*39, 40*). More recently, we identified a potent complement inhibitory activity for the C-terminal domain of BBK32 (BBK32-C, hereafter) (*33*). BBK32-C binds directly to the C1 complex and blocks the earliest proteolytic events of the classical pathway of complement (*33*). The classical pathway-selective inhibitory activity of BBK32 is currently unique among known microbial complement evasion proteins.

We have previously shown that BBK32-C binds the human C1 complex by specifically recognizing the C1r subcomponent with high-affinity (*33*). Recently we obtained a high-resolution crystal structure of BBK32-C and mapped its binding site on human C1r to the serine protease (SP) domain (*41*). Collectively, these studies showed that BBK32-C blocks C1r autoactivation, as well as C1r-mediated cleavage of its partner protease in the C1 complex, C1s (*33*). However, the structural basis and mechanism of action for BBK32-mediated C1r inhibition remains unknown. To gain further insight into the nature of the BBK32/C1r protein-protein interaction, we carried out a series of co-crystallography and size exclusion chromatography-coupled small angle x-ray scattering (SEC-SAXS) experiments followed by a structure-guided mutagenesis strategy involving biophysical, biochemical, and microbiological approaches. These investigations reveal the structural basis for C1r inhibition by Lyme disease spirochetes, provide insight into BBK32’s C1r selectivity, and reveal a convergent inhibitory mechanism between a spirochete and a hematophagous eukaryotic organism.

## Results

### The Structure of BBK32-C in Complex with Human C1r

We have previously shown that BBK32-C interacts with high affinity with an autolytic C-terminal region of human C1r corresponding to residues 300-705 (UNIPROT numbering) (*41*). Co-crystals of BBK32-C in complex with this fragment of C1r produced crystal samples that generally did not yield diffraction data beyond 10 Å limiting resolution. Ultimately, a single sample was produced that diffracted to 4.1 Å. Unfortunately, this dataset was characterized by significant data pathologies including severe anisotropy and low completeness. Despite this, a reasonable final solution (PDB: 7MZT) could be obtained for a single copy of the BBK32-C/C1r complex for which readily interpretable electron density maps were observed (**Fig. 1A, S1 A-C** and **Table S1**). A large protein-protein interface between BBK32-C and the SP domain of C1r, totaling 1,340 Å^2^ buried surface area (b.s.a.), was present in this structure (**Fig. 1A, D, S1 A**). However, underlying data quality issues prevented a complete modeling of the asymmetric unit in this crystal (see *Materials and Methods* for a detailed description). This led to elevated R_free_/R_work_ values (36.9%/37.2%) outside of the normal range for high quality crystallographic solutions (i.e. < 30% R_work_). Despite extensive effort to overcome these issues, including co-crystallization trials with a variety of SP-containing C1r constructs, ultimately an improved crystallographic model could not be obtained. We thus turned to an orthogonal approach involving the study of the BBK32-C in complex with recombinant C1r-CCP2-SP in solution by SEC-SAXS.

**Figure 1.**
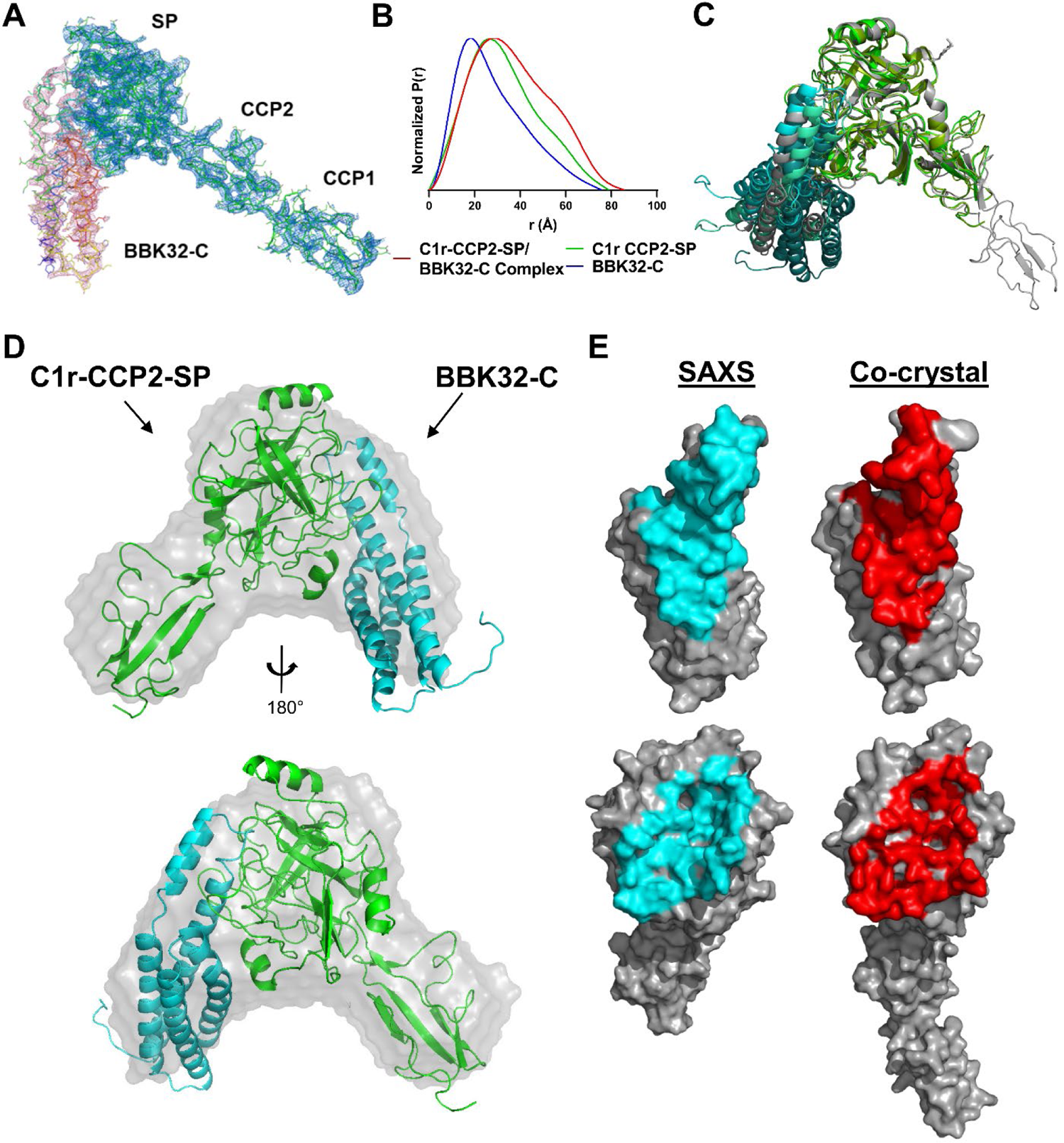
Structures of BBK32-C in complex with human C1r fragments. **A)** The co-crystal structure of BBK32-C in complex with an autolytic fragment of C1r (residues 300-705) at 4.1Ǻ (PDB: 7MZT) is shown with 2*Fo-Fc* density contoured at 1.2 σ for the BBK32-C polypeptide (Chain I; pink mesh) and C1r (Chains A, B; blue mesh). **B)** Normalized pair distribution curves (P(r)) of C1r-CCP2-SP/BBK32-C complex (red), C1r-CCP2-SP (green), and BBK32-C (blue) derived from SEC-SAXS experiments. **C)** An overlay of the top three BBK32-C/C1r-CCP2-SP SAXS models obtained from FoXS Dock as judged by χ^2^ where C1r-CCP2-SP are colored in shades of green and BBK32-C in shades of blue. PDB: 7MZT was aligned to the C1r SP domain of the SAXS structures (grey). The protein-protein interface in each structure is similar with the largest differences being attributed to the orientation of the helical bundle that does not participate in the interface. **D)** *Ab initio* SAXS envelope (gray) aligned with SAXS solution structure from FoXSDock, refined with BILBOMD. **E)** Contact residues as judged by the EMBL PDBePISA server for the BBK32-C/C1r-CCP2-SP SAXS structure (left, cyan) and the BBK32/C1r co-crystal structure (right, red).

**Table 1.** MALS and SAXS parameters

| Sample | Maximal<br>dimension<br>$D_{\max}$ (Å) | $R_g$ (Å) | $MW_{\text{seq}}$<br>(kDa) | $MW_{\text{SAXS}}$<br>(kDa) | $MW_{\text{MALS}}$<br>(kDa) | SAXS<br>model fit<br>$\chi^2$ |
| --- | --- | --- | --- | --- | --- | --- |
| BBK32-C | ~75 | $21.3 \pm 0.9$ | 16.9 | $19.9 \pm 1.58$ | ~19 | 2.08 |
| C1r-CCP2-SP | ~80 | $24.0 \pm 0.9$ | 37.7 | $31.7 \pm 1.75$ | ~40-60 | 1.06 |
| BBK32-C/<br>C1r-CCP2-SP<br>Complex | ~85 | $27.8 \pm 2.2$ | 54.6 | $55.6 \pm 4.13$ | ~55-65 | 1.24 |

SEC-SAXS curves were obtained for recombinant BBK32-C (**Fig. S2A, D**), a recombinant autoactivated C1r domain truncation construct (hereafter, C1r-CCP2-SP) (**Fig. S2B, E**), and an equimolar mixture of BBK32-C and C1r-CCP2-SP (**Figs. S2C, S3**). Radius of gyration (R_g_) values of 21.3, 24.0, and 27.8 Å were calculated from the Guinier region of the scattering curves for BBK32-C, C1r-CCP2-SP, and BBK32-C/C1r-CCP2-SP samples, respectively (**Table 1**, **Figs. S2D, E, S3 insets**). SAXS and multi-angle light scattering (MALS) determined molecular weights (i.e., MW_SAXS_ and MW_MALS_) are consistent with BBK32-C and C1r-CCP2-SP behaving as monomeric species in solution (**Table 1**). For the BBK32-C/C1r-CCP2-SP sample, MW_SAXS_ = 55.6 kDa (*42*) and MW_MALS_ = ∼55-65 kDa were determined, which are close to the expected molecular weight of a 1:1 complex (MW_sequence_ = 54.6 kDa). Analysis of each scattering curve using a pair-wise distribution function, P(r), is also consistent with complex formation as evidenced by the shape of the overlaid P(r) profiles and increased maximum particle size (*D*_max_) of the BBK32-C/C1r-CCP2-SP samples relative to BBK32-C or C1r-CCP2-SP alone (**Fig. 1B, Table 1)**.

Next, we compared the solution state of each unbound protein to available high-resolution crystal structures of BBK32-C (PDB: 6N1L (*41*)) and the activated form of C1r-CCP2-SP (PDB: 1MD8 (*43*)). First, MODELLER (*44*) as implemented in UCSF Chimera (*45*), was used to model missing loops into each structure, and theoretical scattering profiles of each model were then generated and compared against the raw scattering profiles using Fast X-ray Scattering (FoXS) (*46, 47*). Good agreement between theoretical and experimental SAXS curves were obtained for BBK32-C (χ^2^ = 2.08) and C1r-CCP2-SP (χ^2^ = 1.06) atomistic models (**Fig. S2D, E,Table 1**). Collectively, analysis of the experimental SAXS curves shows that solution states of unbound BBK32-C and activated C1r-CCP2-SP are in agreement with previously published crystal structures and that BBK32-C/C1r-CCP2-SP forms a 1:1 stoichiometric complex in solution.

To better define the molecular interface formed between BBK32 and human C1r, we employed a SAXS-restrained macromolecular computational docking approach using FoXSDock (*48*). This method – which relies on the availability of high-resolution structures of the individual complex components – involves initial docking using a geometric shape-matching algorithm, coarse SAXS filtering of the resulting docking models, scoring of the interface energies, and scoring the fit of the model’s theoretical SAXS profiles to the raw SAXS profile of the BBK32-C/C1r-CCP2-SP complex (*47*). BBK32-C and C1r-CCP2-SP were docked as a ligand-receptor pair and each model was compared against the BBK32-C/C1r-CCP2-SP SAXS curve. The top four models (as judged by Z scores (*48*)), each exhibited nearly identical protein-protein interfaces involving BBK32-C contact residues originating from helix 1, 5 and the short loop connecting helix 1 and 2. Consistent with our previous mapping study (*33*), all C1r interactions in these four models involve only residues within the C1r SP domain. We then reconstructed a molecular envelope using DAMMIF (*49, 50*), which produced a distinctive chevron shape (**Figs. 1D,** gray envelope). Superimposition of the top scoring FoxsDock model onto the shape envelope results in a close match, further supporting the validity of the interface (**Fig. 1D**). Minimal molecular dynamics simulations using BILBOMD (*51*) were performed on the best scoring model (χ^2^ = 1.28) to eliminate potential clashes between side chains at the interface, further improving the model’s SAXS score (χ^2^ = 1.24) (**Fig. 1D,Table 1**). A structural alignment of the co-crystal structure to the top three scored SAXS-derived structures reveals a highly similar protein-protein interface, with the primary differences being related to the orientation of the BBK32-C helical bundle that does not participate in C1r interaction (**Fig. 1C, E**). Furthermore, a model of 7MZT that lacks the C1r-CCP1 domain fits closely to the experimental SAXS curve obtained for BBK32-C/C1r-CCP2-SP, as judged by FoXS (χ^2^ = 1.20) (**Fig. S3**).

### Analysis and Validation of the BBK32-C/C1r Structure

Protein-protein interaction analysis was carried out using the SAXS-derived structure of BBK32-C in complex with C1r-CCP2-SP using the programs PISA and CCP4 CONTACT (*52, 53*). The model predicts an interaction involving 28 contact residues on the BBK32-C side and 34 contact residues from human C1r, resulting in of 1,270 Å^2^ b.s.a. (**Fig. 1E, 2A**). To validate the predicted interface, we carried out structure-guided site-directed mutagenesis. A total of eleven BBK32-C residues comprising > 50% of the total surface area buried by BBK32-C were changed to alanine in single and/or double mutant forms (**Fig. 2A**, red). These mutants were then evaluated for direct relative binding to C1r-CCP2-SP by surface plasmon resonance (SPR).

**Figure 2.**
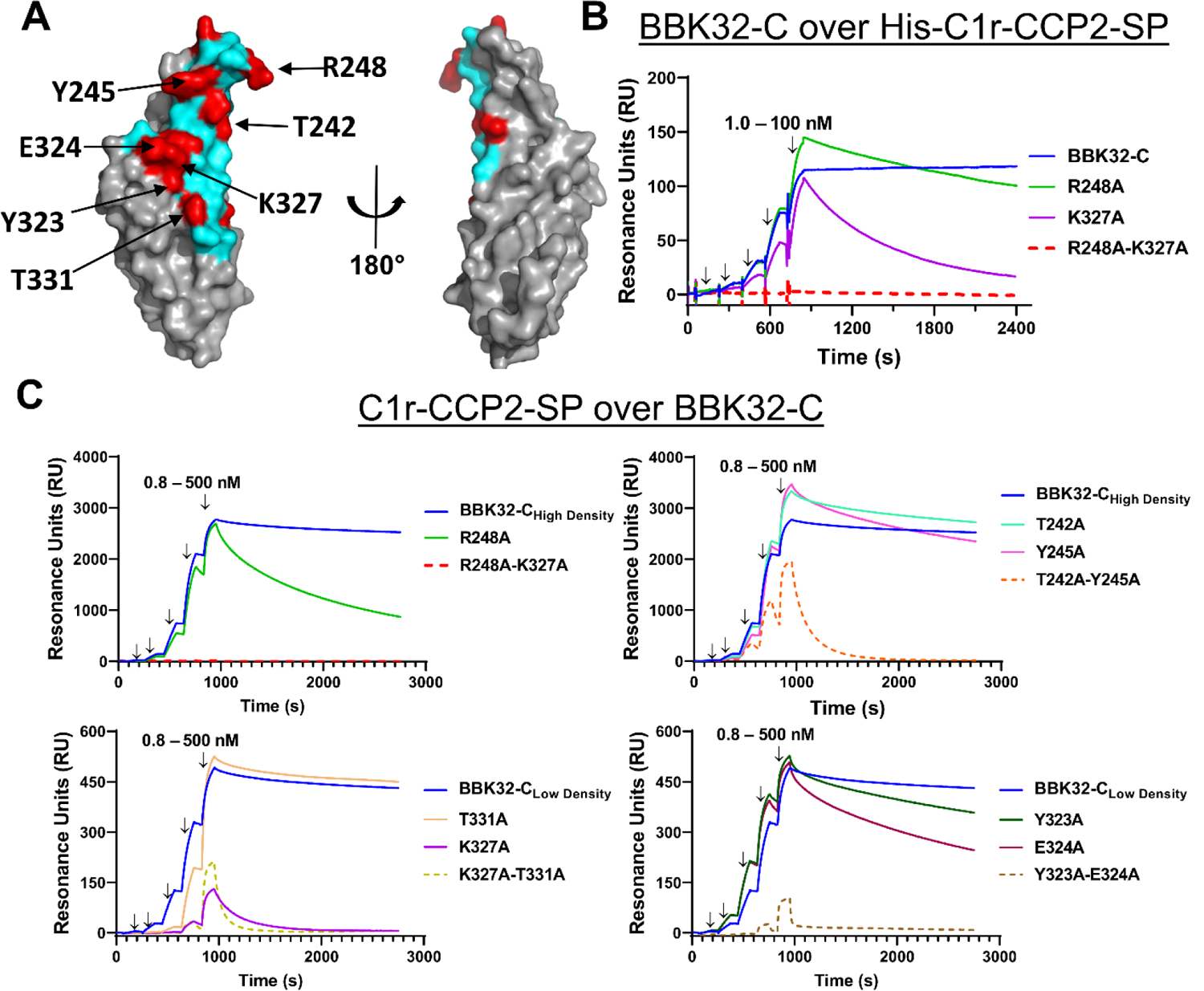
SPR binding assays with structure-guided site-directed BBK32 mutants. **A)** Surface representation of BBK32-C with non-contact surface areas (gray) and BBK32-C/C1r-CCP2-SP contact area (cyan). Residues mutated for this study (red) are indicated in **Table S2**. Site-directed mutants used in Fig. 2B-C are denoted with arrows. **B)** SPR experiments were performed by capturing His-C1r-CCP2-SP on a Ni-NTA sensor chip. BBK32-C and selected site-directed mutants were injected over the His-C1r-CCP2-SP surface in a concentration series (1.0, 3.2, 10.0, 31.6, 100 nM). Injection phases are denoted with arrows. **C)** SPR experiments were performed in the reverse orientation compared to panel **B** by immobilizing each BBK32 protein to sensor chips using amine coupling chemistry. A fivefold concentration series of C1r-CCP2-SP (0.8, 4.0, 20.0, 100, 500 nM) was sequentially injected over each immobilized BBK32-C mutant. Arrows indicate injection phases of each C1r-CCP2-SP concentration. Representative sensorgrams are shown with ligands of similar immobilization density grouped with either high immobilization density BBK32-C (i.e., BBK32-C_High Density_) or low immobilization density BBK32-C (i.e., BBK32-C_Low Density_). BBK32-C curves are reproduced in each respective panel for comparison. SPR experiments were performed in duplicate. Dissociation constants (*K*_D_) were calculated by kinetic fits with a 1:1 Langmuir model (**Fig. S4, S5, Table S2**). Additional site-directed mutants analyzed by SPR and are shown in **Fig. S5**.

To evaluate BBK32 mutants on the same surface, we initially captured HIS-C1r-CCP2-SP uniformly on an Ni-NTA sensorchip and binding of BBK32-C was monitored using a single-cycle injection strategy (*54*) (**Fig. 2B**). Binding of BBK32-C to the HIS-C1r-CCP-2SP biosensor is characterized by a long complex half-life (**Fig. 2B**, blue lines). However, this feature combined with a small upward baseline drift of the reference-subtracted sensorgram, which was related to the affinity-capture-based surface, resulted in poorly determined equilibrium dissociation *K*_D_ values when fitted to a single-cycle kinetic model of binding. To overcome this limitation, and to better match the expected physiological orientation of the interaction under study, we evaluated C1r-CCP2-SP analyte injections over immobilized BBK32 sensorchips generated by amine-coupling chemistry (**Fig. 2C**). Baseline drift was minimal using this covalently-coupled biosensor. BBK32-C biosensors were created at two immobilization densities for better comparison to individual site-directed mutant surface densities. The resulting low and high density sensorgrams fit well to a 1:1 Langmuir kinetic binding model as judged by residual plots (**Fig. 2C** blue lines, **Figs. S4**, **S5,** and **Table S2**). Consistent with our previous SPR binding measurements of BBK32-C for C1, C1r, an autolytic fragment of C1r, and HIS-tagged recombinant domain truncations of C1r (*33, 41*), BBK32-C bound to activated C1r-CCP2-SP with high affinity, exhibiting an equilibrium dissociation constant (*K*_D_) of 0.45 nM (high density surface) and 0.78 nM for the low density surface (**Fig. 2C,Table S2**).

Each BBK32 site-directed mutant was then evaluated by generating individual biosensors, carrying out a C1r-CCP2-SP titration series, calculating an associated *K*_D_, and comparing to the BBK32-C biosensor that most closely matched wild-type surface density (**Fig. 2C, S4, S5**). Three different double alanine mutants involving mutagenesis of small contiguous contact surfaces resulted in a 653-fold (K327A-T331A), 127-fold (T242A-Y245A), and 615-fold (Y323A-E324A) reduction in affinity for C1r-CCP2-SP, whereas a fourth such mutant retained an affinity similar to wild-type (L236A-T239A) (**Figs. 2C, S4, S5, Table S2**). In the case of T242A-Y245A and Y323A-E324A, single mutants retained wild-type-like affinity, indicating that the observed affinity loss in the double mutants was due to a cumulative disruption of a larger contact surface area (**Fig. 2C**). In contrast, a single K327A mutant resulted in a 153-fold loss in affinity, whereas the paired T331A single mutant had near wild-type affinity (**Fig. 2C**). Additional mutagenesis was directed at two arginine residues (R248 and R337). While a single R337A mutant exhibited wild-type like affinity, mutation of R248 resulted in a 13-fold loss in affinity for C1r-CCP2-SP (**Fig. 2B, C, S4, S5**). Although they do not form a contiguous binding surface, a pairing of the two strongest loss-of-affinity single mutants (i.e., R248A-K327A), produced a double mutant where no detectable binding signal was observed (**Fig. 2B, C, S5**). In all cases, loss of affinity was driven by increased dissociation rate constants (*k*_d_) (**Table S2**). Similar qualitative results were obtained for the single and double mutants of R248A and K327A as analytes over the HIS-C1r-CCP2-SP biosensor (**Fig. 2B**), indicating that surface orientation and/or surface density did not greatly influence the measurement of relative binding strength for these mutants. Taken together, these data show that many BBK32 residues predicted to contact C1r-CCP2-SP in the SAXS structure (**Fig. 2A**) exhibit a loss of affinity when mutated to alanine and that R248 and K327 are critical hotspot residues for mediating C1r-CCP2-SP interaction.

### BBK32 Occludes Subsites Within the Active Site of C1r

Having taken steps to validate the structural models, and in so doing, identifying key interactions that mediate BBK32-C/C1r complex formation, we then analyzed the structure for potential clues into the mechanism of C1r inhibitory action by BBK32. C1r is a modular serine protease with a chymotrypsin-like SP domain (*55*). We noted that the BBK32 binding site on human C1r is in close proximity to the C1r catalytic site (**Fig. 3A**). Although limited contacts are made in the model with active site residues (28 Å^2^ b.s.a.), much more extensive interactions are made with the specificity pocket subsites S1 (176 Å^2^ b.s.a.) and S1’ (167 Å^2^ b.s.a.) (**Fig. 3B**). Residues involved in S1 (red) and S1’ (blue) subsite interactions were mapped onto the BBK32-C surface (**Fig. 3B**). These include three residues (T242, Y245, and R248) that were shown to have loss of affinity effects when mutated to alanine (**Fig. 2**).

**Figure 3.**
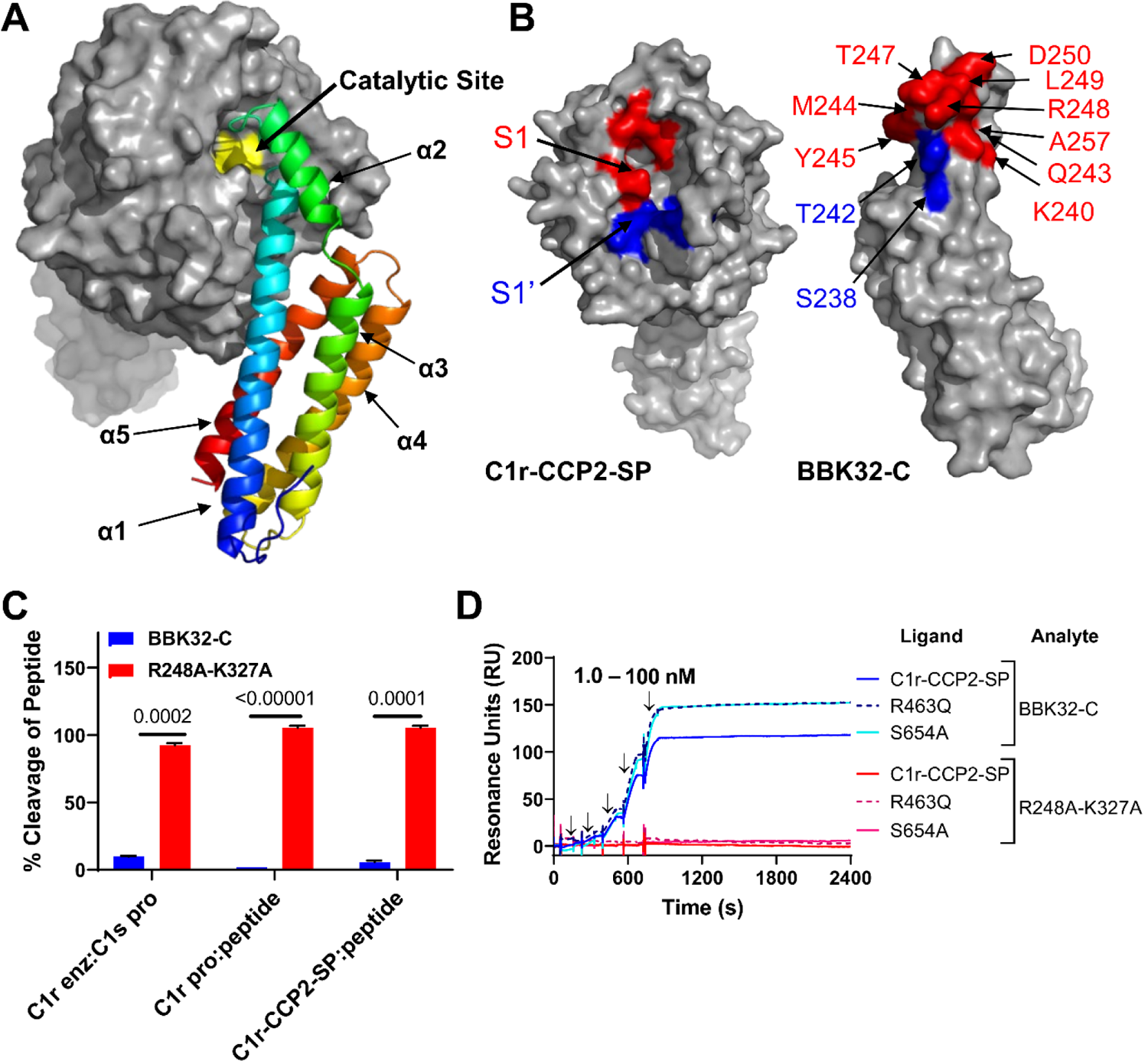
BBK32-C occludes the S1 and S1’ subsites on C1r. **A-B)** C1r-CCP2-SP shown with the catalytic site (yellow), substrate specificity subsites S1 (red) and S1’ (blue) as determined by Budayova and colleagues (*43*) using nomenclature provided by Schechter and Berger (*88*). BBK32-C shown with residues that contact the C1r S1 site (red) and S1’ site (blue). **C)** The ability of BBK32-C or BBK32-R248A-K327A to directly inhibit cleavage of a small peptide or a natural substrate was evaluated by colorimetric substrate-based enzyme assays. BBK32-C significantly blocked C1r enzyme cleavage of C1s proenzyme, autocatalysis of C1r proenzyme and C1r-CCP2-SP enzyme. All experiments were performed in triplicate and normalized to positive (no inhibitor) and negative (no enzyme) controls. Statistical significance was evaluated using paired t-tests with p-values noted above each experiment. **D)** To evaluate BBK32-C binding to zymogen-stabilized forms of C1r-CCP2-SP, SPR experiments were performed by capturing His-C1r-CCP2-SP, His-C1r-CCP2-SP-R463Q (R463Q), or His-C1r-CCP2-SP-S654A (S654A) on a Ni-NTA sensor chip. BBK32-C was sequentially injected over each surface in a concentration series (1.0, 3.2, 10.0, 31.6, 100 nM). BBK32-C sensorgrams are reproduced from Fig. 2B for comparison. Arrows indicate injection phases of each C1r-CCP2-SP concentration.

The steric occlusion of the S1 and S1’ site by BBK32 in the model predicts that it would also be capable of directly blocking the hydrolysis of small peptide C1r substrates, as well as the larger natural C1r substrates, C1r and C1s proenzymes. Previously, we have shown that BBK32 blocks cleavage of proenzyme C1s by isolated C1r enzyme using an SDS-PAGE-based readout (*33*). We adapted this assay to one that utilizes a colorimetric substrate specific for C1s enzyme (i.e. Z-L-Lys thiobenzyl), and found that BBK32-C, but not R248A-K327A, could prevent activation of proenzyme C1s by C1r enzyme. Similar results were obtained when C1r-CCP2-SP cleavage of a colorimetric peptide substrate that requires only P2-P1-P1’ access (i.e. Z-Gly-Arg-thiobenzyl) was monitored (**Fig. 3C**). Furthermore, BBK32-C, but not R248A-K327A, prevented cleavage of this substrate by spontaneously autoactivated full-length proenzyme C1r (**Fig. 3C**). To further evaluate the interaction of BBK32-C with the proenzyme form of C1r, we produced two forms of zymogen-stabilized C1r-CCP2-SP mutants. Both the scissile loop (i.e. R463Q) and catalytic serine-based (S654A) zymogen forms of C1r-CCP2-SP bound similarly to BBK32-C, and like autoactivated C1r-CCP2-SP, exhibited slow dissociation rates (**Fig. 3D**). Together, these data show that BBK32 directly blocks both small and large C1r substrates from proteolytic cleavage and that BBK32 recognizes both proenzyme and activated enzyme forms of C1r.

### R248 and K327 are Critical for BBK32-C Complement Inhibitory Activity

Next we tested our panel of structure-based recombinant mutants for their ability to block serum-based human complement activation in an ELISA assay that detects the classical pathway activation product C4b. BBK32-C exhibited a half-maximal inhibitory concentration (IC_50_) of 15 nM in this assay format (**Fig. 4A, Table S2**), consistent with our previously reported value of 5.6 nM (*33*). With the exception of E324A, which is discussed below, near wild-type IC_50_ values were measured for all BBK32-C mutants that retained high affinity for C1r-CCP2-SP, although we noted a modest reduction in inhibitory activity in the Y245A (∼5-fold), N251A (∼6-fold) and Y323A (∼2-fold) single mutants (**Fig. S6 and Table S2**). Importantly, in all cases where mutants exhibited a strong loss of affinity for C1r-CCP2-SP, they were also greatly impaired in their ability to block classical pathway activation. For example, the contiguous surface double mutants Y323A-E324A and K327A-T331A failed to block complement at any concentration, while a ∼240-fold weaker IC_50_ value for T242A-Y245A was measured (**Fig. S6 and Table S2**). Likewise, the R248A mutant was attenuated ∼28-fold, and neither the K327A nor the R248A-K327A mutant blocked at any of the concentrations tested (**Figs. 4A, and Table S2**).

**Figure 4.**
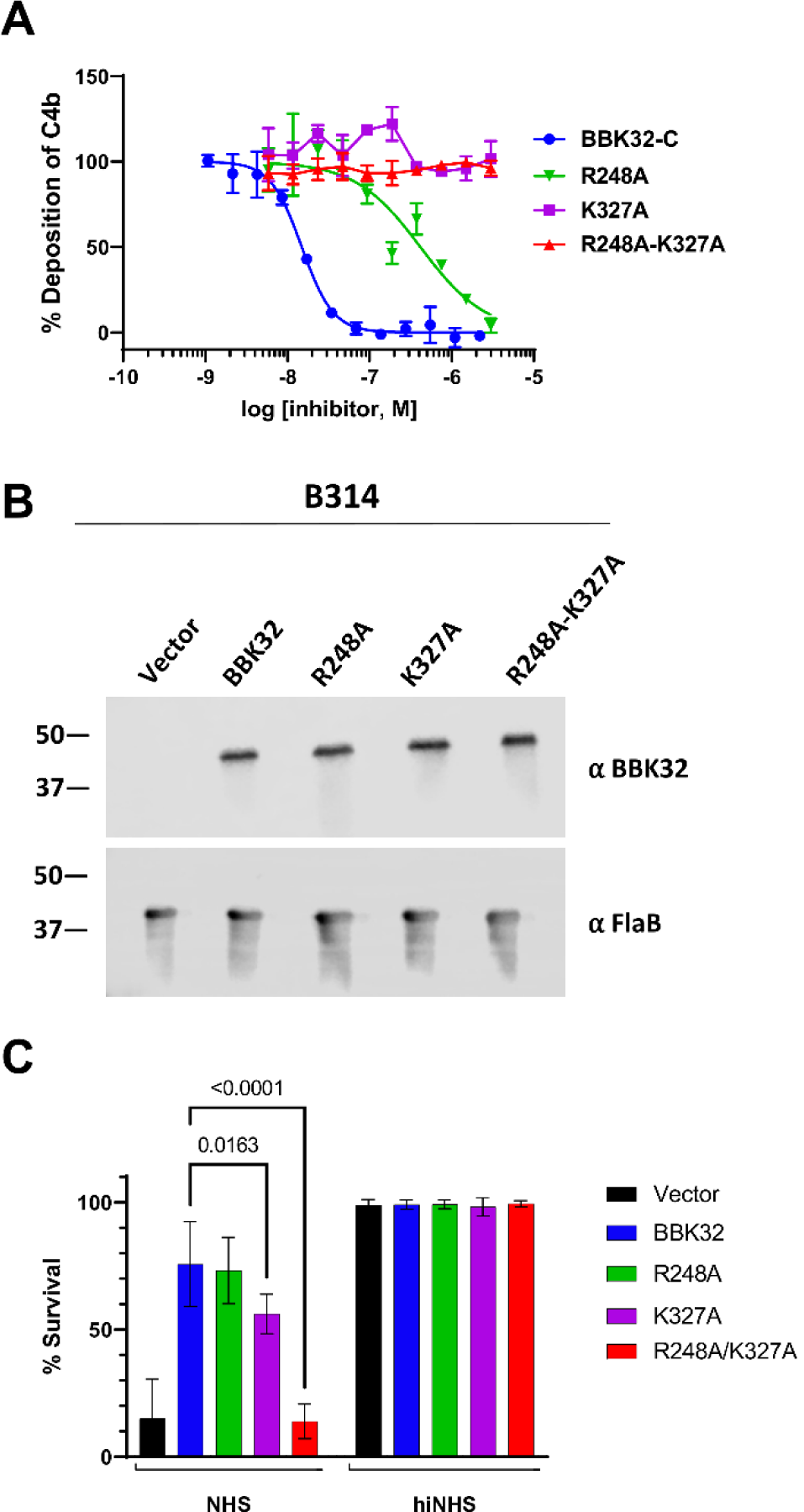
BBK32 R248 and K327 are critical for complement inhibitory activity of BBK32. **A)** ELISA-based complement activation assays were performed to determine the ability of BBK32-C mutants to inhibit the classical pathway of complement *in vitro.* A two-fold concentration series of each BBK32-C mutant was used and performed in duplicate. Inhibitory data for additional site-directed mutants are presented in **Figure S6**. IC**_50_** values and 95% confidence intervals associated with a normalized four-parameter dose-inhibition fitting procedure are shown in **Table S2**. **B)** Western blots of lysates from *B. burgdorferi* strains B314 containing distinct *bbk32* alleles were probed with monoclonal antibodies to BBK32 (α-BBK32) or FlaB (a-FlaB), the latter as a loading control. Vector refers to the plasmid only backbone sample (B314/pBBE22*luc*), BBK32 is the wildtype *bbk32* construct (B314/pCD100), R248A is the *bbk32*-R248A isolate (B314/pAP5), K327A is the *bbk32*-K327A isolate (B314/pAP6), and R248A-K327A is the *bbk32*-R248A-K327A double mutant (B314/pAP7). Numbers on the left refer to the molecular mass of markers (in kDa). **C)** BBK32 mutant proteins were tested for their ability to confer resistance to normal human serum (NHS). The serum sensitive strain B314 was transformed separately with vector only (black), wildtype *bbk32* (blue), *bbk32* R248A (green), *bbk32* K327A (purple), and *bbk32* R248A-K327A (red) and incubated with NHS. Sensitivity was scored as a ratio of the affected cells relative to the total cells viewed via darkfield microscopy. Heat inactivated NHS (hiNHS) was used as a control (right). P-values between *bbk32* samples are indicated above. Statistical significance was assessed using a two-way ANOVA.

Based on the SAXS-derived BBK32-C/C1r-CCP2-SP structure we were able to identify several BBK32 residues that are important for its activity. In particular, substitution of R248 and K327 to alanine in single mutants, or as a double mutant, results in a significant reduction of both C1r-binding and inhibitory activity. To study these residues in a more physiological setting, we produced site-directed BBK32 ‘knock-in’ strains in the *B. burgdorferi* B314 genetic background (**Fig. 4B**). *B. burgdorferi* B314 lacks linear plasmids that encode many of the outer surface proteins associated with infection, including BBK32 (*56, 57*). Previously we have shown that when wild-type BBK32 is expressed heterologously on the surface of *B. burgdorferi* B314, it protects these normally serum-sensitive spirochetes from human complement-mediated bacteriolysis (*33, 41*).

Introduction of the R248A mutant into *B. burgdorferi* B314 resulted in a minimal reduction in inhibitory activity relative to wild-type BBK32, while spirochetes expressing K327A were significantly more sensitive to human serum. Spirochetes transformed with the *bbk32* R248A-K327A allele lost all protective activity and were indistinguishable from the vector-only control (**Fig. 4C**). These data confirmed that the R248 and K327 residues are critical for eliciting complement inhibitory activity in the context of full-length BBK32 on the outer surface of *B. burgdorferi*. Western immunoblots showed that full length BBK32 was produced equivalently in all mutants tested, indicating that the enhanced sensitivity of the R248-K327A double mutant was not the result of proteolytic instability (**Fig. 4B**).

### BBK32-C Targets Non-Conserved C1r Loops Outside of the Active Site

We have previously shown that BBK32 is highly selective for C1r-binding relative to C1s (*33*). To gain insight into this phenomenon, and with no comparable C1r inhibitors having been reported, we instead compared the BBK32-C/C1r-CCP2-SP SAXS structure to that of a previously published crystal structure of a protein inhibitor of both the classical and lectin complement pathways, known as gigastasin (*58*). Gigastasin differs in selectivity from BBK32 as it blocks C1s, MASP-1 and MASP-2, but is a relatively weak inhibitor of C1r (*58*). Interestingly, the gigastasin/C1s protein-protein interface shares common features with the BBK32-C/C1r-CCP2-SP SAXS structure presented here. Namely, both inhibitors form extensive interactions with the S1 and S1’ subsites of their respective target enzymes. Furthermore, BBK32 and gigastasin each interact with three or more residues in the A and D loops, as well as loop 2 (**Fig. 5**). However, BBK32 differs from gigastasin in that it makes extensive interactions with the B loop of C1r. This loop contains an insertion sequence making it longer in C1r compared to most other complement serine proteases, including C1s (**Fig. 5**). Interestingly, the B loop also is thought to block the binding of the antistasin domain of gigastasin making it a determinant of inhibitor specificity (*43, 58*). Of the 16 residues that comprise the C1r B loop, 10 form contacts with BBK32-C and none are identical at the homologous position in C1s (**Fig. 5**). Considering the entire interface, 25 of the 34 C1r contact residues predicted by the BBK32-C/C1r-CCP2-SP SAXS structure are non-identical in C1s (**Fig. 5**). Most of these arise from surfaces involving the S1’ site and four C1r loops (i.e. A, B, D, E loops), suggesting these interactions may underlie the selectivity of BBK32 for C1r. Given the vastly different three-dimensional structures of BBK32 and gigastasin (**Fig. 5**), it appears that a bloodborne prokaryote (*B. burgdorferi*) and a blood-feeding eukaryote (*Haementeria ghilianii*) have evolved complement inhibitors with at least partially convergent mechanisms of action, while also evolving differential target selectivity.

**Figure 5.**
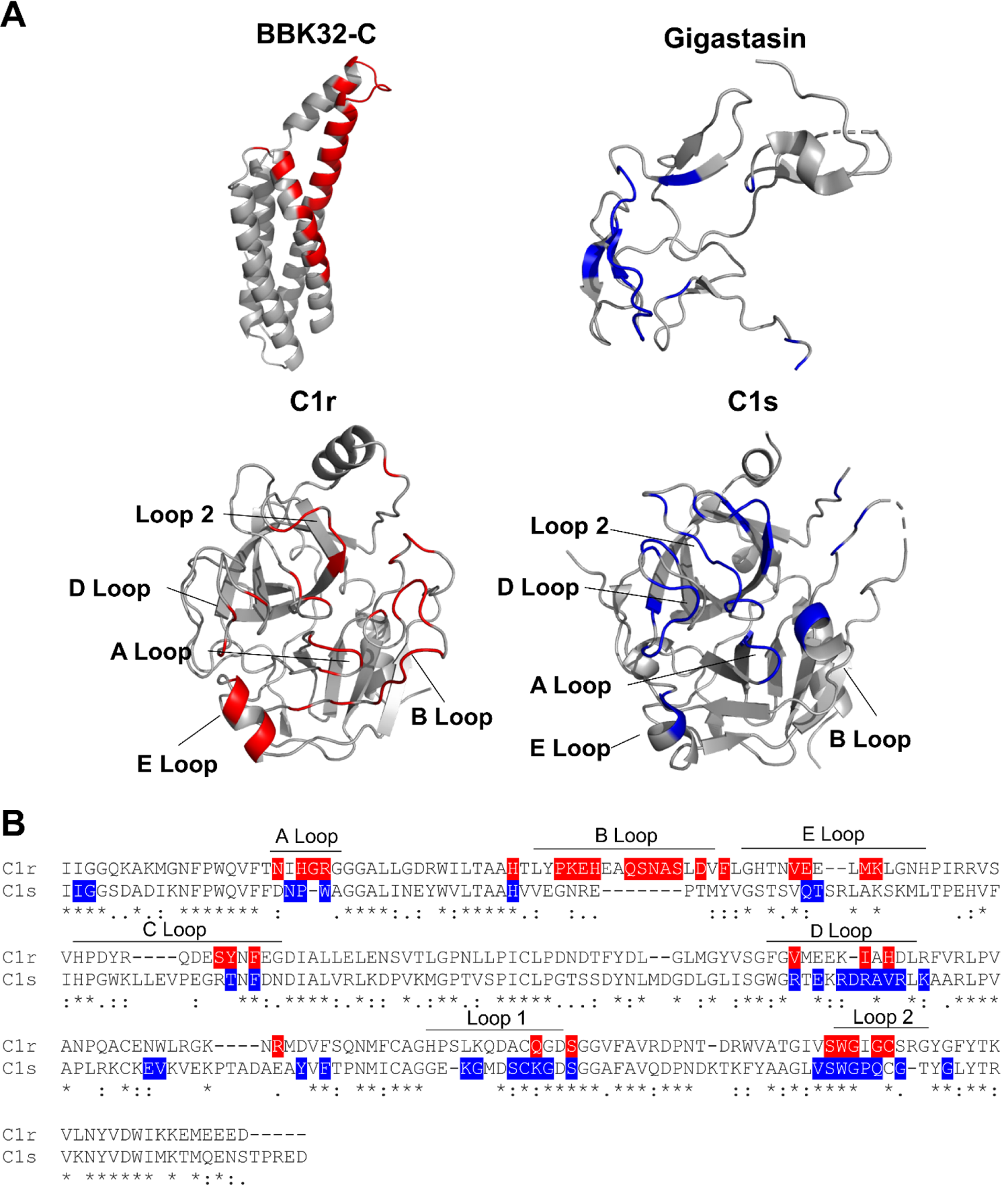
Comparison of BBK32 to gigastasin reveals convergent mechanisms of action. **A)** Ribbon diagrams of BBK32-C (PDB:6N1L), C1r-CCP2-SP (PDB:1MD8), Gigastasin/C1s-CCP2-SP (PDB: 5UBM) with colored interactions within 4 Å, as judged by the program CCP4i CONTACTS. The BBK32-C/C1r-CCP2-SP interactions are shown highlighted in red and C1s/Gigastasin loop interactions in blue. **B)** Sequence alignment of serine protease domains of C1r and C1s loops are designated with bars and are according to C1r structure.

## Discussion

In the absence of regulation, complement activation at the microbial surface leads to deposition of C3b or C4b on the microbe surface, recognition by innate immune cells due to the presence of immobilized C3b and C4b, recruitment of leukocytes by the anaphylatoxins C3a and C5a, stimulation of the adaptive immune response by each of these fragments, and direct bactericidal action by C5b-9 (i.e., MAC) (*1, 2*). While intracellular pathogens may be protected by complement within an intracellular niche, extracellular pathogens like *Borreliella* spirochetes face a continuous threat from the complement systems of their hosts.

*Borreliella* first encounter complement during the bloodmeal of an infected *Ixodes* sp. tick. Following transmission from their arthropod vector into the vertebrate host, spirochetes must continue to protect themselves from complement during hematogenous dissemination, a challenge that persists once they colonize distant tissues. To evade complement attack, Lyme disease spirochetes express nearly a dozen different outer surface lipoproteins that directly interact with complement proteins and modulate their native activities (*16, 17*). While we have previously shown that *B. burgdorferi bbk32* is required for optimal infectivity in naïve mice, it is likely that C1r-specific inhibition by BBK32 also protects *B. burgdorferi* from targeted antibody-dependent complement-mediated killing, a hallmark of the classical pathway. This contention is supported by the persistent infection observed despite the presence of high titered antibody responses against *B. burgdorferi* that can passively protect against borrelial infection and kill *B. burgdorferi in vitro* in a complement-dependent manner (*59–61*).

In this study we aimed to determine the structural basis for the inhibition of C1r by *B. burgdorferi* BBK32. A clear limitation of our study is the low resolution inherent to SAXS experimental data. Importantly, the model presented here assumes limited conformational rearrangement of BBK32-C and C1r-CCP2-SP relative to their unbound forms as previously determined by crystallography. While analysis of the SAXS data for BBK32-C and C1r-CCP2-SP suggests that each unbound protein adopts a similar structure in solution as they do in their respective crystal structures (**Fig. S2**), we recognize that our approach would be unable to detect larger scale conformational changes induced by BBK32-C/C1r-CCP2-SP complex formation. We note that a highly similar interface is observed in the co-crystal structure presented here (**Fig. 1**), although we acknowledge that the high R values and associated issues in the crystallographic data quality are an additional limitation. Nonetheless, the validity of the SEC-SAXS-derived structure is supported by its ability to successfully guide the identification of many BBK32 residues important for driving both affinity and inhibitory activity (**Figs. 2-4**). Furthermore, the structure provides a basis for an inhibitory mechanism that is consistent with the inhibitory activities and selectivity of BBK32 (**Figs. 3-5**). Taken together, our data suggests that BBK32 inhibits human C1r by simultaneously targeting its active site, as well as functionally critical, less conserved, non-active site residues on loops of the C1r SP domain.

Like Lyme disease spirochetes, other bloodborne pathogens and blood feeding eukaryotes have evolved complement evasion systems. For example, potent direct inhibitors of complement are produced in the salivary glands of several species of ticks, including *Ixodes* sp. (*62–66*). Similarly, blood-feeding insects like sandflies (*67*) and mosquitos (*68*), along with other human extracellular pathogens such as *Staphylococcus aureus* (*15, 69, 70*), all produce a wide array of mechanistically distinct protein inhibitors of complement. However, among known complement evasion molecules, relatively few target the classical pathway with specificity (*70*), and to our knowledge BBK32 remains the only reported selective inhibitor of C1r. Despite this, our study identifies a surprising convergence of function between *B. burgdorferi* BBK32 and a complement protease inhibitor of the classical and lectin pathways previously identified in the giant Amazonian leech (i.e. gigastasin). However, gigastasin is related to a class of inhibitors known as antistasins (*71*), that preferentially bind to the active form of their enzyme targets. This is in contrast to BBK32, which recognizes both proenzyme and enzyme forms of C1r (**Figs. 2, 3**). While each inhibitor functions, at least in part, by steric occlusion of the subsites upon enzyme-inhibitor complex formation (*58*), it appears that differences in non-active site loops may drive selectivity of these inhibitors (**Fig. 5**).

*Borreliella* spirochetes vary widely in their susceptibility to serum-based complement-mediated bacteriolysis (*72*). While the categorizations are general and not without exceptions, *B. burgdorferi* is typically considered serum-resistant whereas *B. garinii* is susceptible to killing by human serum. The overall susceptibility of a spirochete to a particular host complement system is related to the collective activities of the larger *Borreliella* complement evasion system (i.e., CRASPs, OspC, BBK32, etc.). However, insight into this host association phenomenon may be gained from analysis of a single *Borreliella* complement inhibitor, in this case BBK32. While the C1r-CCP2-SP binding affinity of nearly all mutants in our study correlated with the relative inhibitory activity, a curious exception was BBK32-E324A. This mutant was reduced only two-fold in affinity for C1r-CCP2-SP (**Fig. 2, S4**), but by several orders of magnitude in inhibitory activity (**Fig. S6, Table S2**). BBK32-E324 is located next to K327 on the face of alpha helix 5 of BBK32 and, like K327, is positioned across from the B loop of C1r in the model. In our previous study we coincidentally probed the function of E324 while investigating the reduced complement inhibitory activity of a BBK32 homologue from *B. garinii*, termed BGD19 (*41*). In that study, a triple mutant chimeric recombinant protein termed BXK32-C was created to convert BBK32 to BGD19 at three non-conserved residue positions (i.e., BBK32-E308K-Q319K-E324Q) (*41*). This mutant shifted from BBK32-like to BGD19-like inhibitory activity, leading us to conclude that at least one of these three surface-exposed residues was important for the complement inhibitory activity of BBK32. Unlike R248 and K327, which drive human C1r affinity and are universally conserved across publicly available *B. burgdorferi* genome sequences, position 324 is less conserved (**Fig. S7**). While the underlying reasons for the disconnect between the binding and inhibitory activities of these proteins remain unclear, we speculate that variation in position 324 in naturally occurring BBK32 sequences may allow for optimal inhibition of C1r in other vertebrates as the sequence of the C1r B loop is one of the least conserved features within vertebrate C1r. Further studies, including high-resolution structural information involving synthetic or naturally occurring BBK32 sequences such as *B. garinii* BGD19, along with mechanistic studies involving non-human C1r or extensive C1r mutagenesis, will be needed to further test this possibility.

For the first time, this study provides a structural basis for classical pathway complement evasion by a human pathogen. Among naturally occurring complement inhibitors, *B. burgdorferi* BBK32 is unique in its ability to selectively target and inhibit the initiator protease of the classical pathway, C1r, and, as such, the structure-function studies reported here provide new insight into an evolutionarily optimized mechanism of action for C1r inhibition. This knowledge improves our understanding of complement evasion by bacterial pathogens, increases our understanding of naturally occurring complement inhibitors, and provides a potential basis for the development of novel complement directed therapeutics that are classical pathway-specific.

## Materials and Methods

### Strains, Plasmids constructed, and Oligonucleotides

All bacterial strains, plasmid constructs, and oligonucleotides used are listed in supplemental **Table S3**. *B. burgdorferi* strain B314, with all recombinant plasmid constructs, was grown in complete BSK-II media containing 300 µg/ml kanamycin as previously described (*41*). *E. coli* cells were grown in LB media with kanamycin at 50 µg/ml.

Plasmid constructs for usage in *B. burgdorferi* strain B314 were constructed in a step-wise manner. First, pCD100 was used as template to obtain the *bbk32*-R248A allele through use of the oligonucleotides R248A mutant F and R248A mutant R that face in the opposite orientation and contain the sequence to change codon 248. To facilitate this, GXL polymerase (Takara Bio) was used with the following parameters: a 15 second denaturing step at 98°C, an annealing temperature dependent on the lowest melting temperature of primers used for a specific reaction, and a 68°C extension step for 1min/kb. The reaction was then digested with *Dpn*I (NEB), to digest methylated parental DNA, and transformed into NEB5-alpha competent *E. coli*. The resulting plasmid was designated pAP5. The *bbk32*-K327A encoding construct was obtained by synthesizing a 241 bp fragment starting at nucleotide 861 and encompassing the remainder of *bbk32* and a 40 base pair region of downstream sequence (IDT gBlocks). This fragment of *bbk32* contains native sequence with the exception of the K327A codon. The K327A containing fragment and either pCD100 (for pAP6 construction) or pAP5 (for pAP7 construction) were used as template with oligonucleotide primers described in **Table S3** for PCR amplification. The resulting PCR products were assembled using the NEBuilder HiFi DNA Assembly Cloning Kit as described in (*41*). The construct encoding the *bbk32*-K327A allele was named pAP6. The final construct that encoded the *bbk32*-R248A/K327A double mutant was designated pAP7. All plasmids were sequenced to ensure that the desired mutations were obtained and that no additional mutations were introduced into *bbk32*. Transformations of strain B314 with the aforementioned plasmid constructs were done as described in (*73, 74*).

For the production of all recombinant BBK32 proteins, DNA fragments for corresponding site directed mutations in BBK32-C (residues 206-348, *B. burgdorferi* B31), for expression in *E. coli*, were *E. coli* codon optimized and synthesized commercially by IDT Technologies gBlock Gene Fragment service. The R337A mutant contained the C-terminal 6 residues in the full length BBK32 sequence making it residues 206-354 (*33*). IDT gBlocks were designed with 5’ BamHI, 3’ NotI sites and stop codon, which then were subsequently cloned into pT7HMT as previously described (*33, 75*). Wild-type BBK32-C and C1r-CCP2-SP were expressed and purified form previously generated constructs (*33, 41, 75*).

### Proteins

Recombinant BBK32-C proteins were expressed and purified as previously described (*33, 41*), with the following modifications. After elution in Ni-NTA elution buffer (20 mM Tris (pH 8.0), 500 mM NaCl, 500 mM imidazole (pH 8.0)), proteins were exchanged into Ni-NTA-binding buffer (20 mM Tris (pH 8.0), 500 mM NaCl, 10 mM Imidazole (pH 8.0)) using a Hi-Prep Desalting 26/10 column (GE Healthcare). To remove the HIS-myc affinity tag, proteins were incubated with HIS tagged tobacco etch virus (TEV) protease and 5mM β-mercaptoethanol at room temperature overnight. The digested proteins were then passed over a 5mL HisTrap-FF column and the flow-through was collected for further purification by gel filtration chromatography using a HiLoad Superdex 75 26/600 pg column (GE Healthcare). The monodisperse peak corresponding to BBK32-C was pooled following analysis of chromatography fractions by SDS-PAGE and exchanged into HBS (10mM HEPES (pH 7.3), 140mM NaCl). C1r-CCP2-SP, C1r-CCP2-SP zymogen mutants, and His-C1r-CCP2-SP were purified according to methods as described previously, with a final polishing step involving gel filtration chromatography on a HiLoad Superdex 75 column (GE Healthcare) (*41, 55, 76*). His-tagged proteins were not subjected to HIS-TEV cleavage. An autoproteolytic fragment of human C1r corresponding to residues 300-705 (UNIPROT: P00736) used in the co-crystallography experiment was obtained as previously described (*41*). Full length complement proteins C1r proenzyme, C1r enzyme, and C1s proenzyme were purchased from Complement Technology. In order to test for global changes in protein folding that may be induced by site-directed mutagenesis, recombinant proteins were evaluated for major changes in size by size exclusion chromatography (**Fig. S8**). Proteins were separated on a HiLoad Superdex 200 Increase 10/300 column (GE Healthcare) and plotted against BioRad Standards to determine size. Monodisperse peaks for all BBK32-C related samples exhibited a ranging between 13.2 and 16.4 kDa.

### Statistical analysis

Statistical analysis of assays was performed on GraphPad version 8. Calculations of IC_50_ for classical pathway ELISA was determined with a normalized variable four-parameter fit. Statistical significance for colorimetric enzyme assays were analyzed using a paired t-test. Significance for *B. burgdorferi* serum sensitivity assays were analyzed using a two-way ANOVA with a Šidák correction for multiple comparisons.

### Size exclusion chromatography-coupled Small Angle X-ray Scattering Data Collection

Volumes of 60 µl samples containing either 8.0 mg mL^-1^ BBK32-C, 2.0 mg mL^-1^ C1r-CCP2-SP, or 2.0 mg mL^-1^ BBK32-C/C1r-CCP2-SP 1:1 Complex were prepared in 10 mM HEPES pH 7.3, 140 mM NaCl buffer. SEC-SAXS-MALS were collected at the ALS beamline 12.3.1 LBNL, Berkeley, California (*77*). X-ray wavelength was set at λ=1.127 Å and the sample-to-detector distance was 2100 mm resulting in scattering vectors, q, ranging from 0.01 Å^-1^ to 0.4 Å^-1^. The scattering vector is defined as q = 4πsinθ/λ, where 2θ is the scattering angle. All experiments were performed at 20°C and data was processed as described (*78*). Briefly, a SAXS flow cell was directly coupled with an online Agilent 1260 Infinity HPLC system using a Shodex KW802.5 column. The column was equilibrated with running buffer (10 mM HEPES (pH 7.3), 140 mM NaCl) with a flow rate of 0.5 mL min^-1^. 55 µL of each sample was run through the SEC and three second X-ray exposures were collected continuously during a 30-minute elution. The SAXS frames recorded prior to the protein elution peak were used to subtract all other frames. The subtracted frames were investigated by radius of gyration (Rg) derived by the Guinier approximation I(q) = I(0) exp(-q^2^Rg2/3) with the limits qRg<1.5. The elution peak was mapped by comparing integral of ratios to background and Rg relative to the recorded frame using the program SCÅTTER. Uniform Rg values across an elution peak represent a homogenous assembly (**Fig. S2**). Final merged SAXS profiles, derived by integrating multiple frames across the elution peak, where used in PRIMUS (*79*) for further analysis including Guinier plot which determined aggregation free state (**Figs. S2D-E, S3**). The program GNOM (*80*) was used to compute the pair distribution function (P(r)) (**Fig. 1B**). The distance r where P(r) approaches zero intensity identifies the maximal dimension of the macromolecule (D_max_). P(r) functions were normalized to peak maxima. Eluent was measured in line with a series of UV at 280 and 260 nm, multi-angle light scattering (MALS), quasi-elastic light scattering (QELS), and refractometer detector. MALS experiments were performed using an 18-angle DAWN HELEOS II light scattering detector connected in tandem to an Optilab refractive index concentration detector (Wyatt Technology). System normalization and calibration was performed with bovine serum albumin using a 45 μL sample at 10 mg mL-1 in the same SEC running buffer and a dn/dc value of 0.19. The light scattering experiments were used to perform analytical scale chromatographic separations for MW_MALS_ determination of the principle peaks in the SEC analysis (**Fig. S2A-C**). UV, MALS, and differential refractive index data was analyzed using Wyatt Astra 7 software to monitor the homogeneity of the sample across the elution peak complementary to the above-mentioned SEC-SAXS signal validation (**Fig. S2**).

### SAXS Solution Structure Modeling

High resolution crystal structures for BBK32-C (PDB:6N1L) and C1r-CCP2-SP (PDB:1MD8) were used for structural modeling (*41, 43*). Missing residues, including the GSTGS sequence cloning artifact, were added using COOT (*81*). In order to test multiple configurations of residues added in this manner, structures were imported into UCSF-Chimera and the MODELLER (*82*) tool was used to make five models. Each model was tested against SAXS data in FoXS and the top scoring model was accepted. The BBK32-C/C1r-CCP2-SP complex was determined by using FoXSDock (*47*) to perform rigid docking the highest overall score of the almost 5000 models were accepted. Minimal molecular dynamics (MD) simulations using BILBOMD (*51*) was performed on the best scoring model to eliminate potential clashes between the residues side chains at the interface. The experimental SAXS profiles were then compared to theoretical scattering curves generated from molecular models using the FOXS (*46, 48*). Shape envelope reconstruction for the BBK32-C/C1r-CCP2-SP sample was carried out using DAMMIF as implemented in the ATSAS software suite (*49, 83*). The final model was produced from the average of ten models filtered for low occupancy using default values for DAMAVER/DAMFILT (*50*).

### Crystallization, Structure Determination, Refinement and Analysis

Purified BBK32-C and a purified autolytic fragment of human C1r (Complement Technologies) were mixed at 5% molar excess BBK32-C and concentrated to 3.5 mg ml^-1^ in 10 mM HEPES (pH 7.3) 140 mM NaCl. Small plate-like crystals were obtained by vapor diffusion in hanging drops at 20 °C by mixing 1 µl of protein solution with 1 µl of precipitant solution that had been diluted into 3 µl of double deionized water. The precipitant solution contained 0.2 M sodium formate and 27.5% (v/v) PEG 3,350. Crystals appeared in 2-5 days and were harvested and cryoprotected in the precipitant solution supplemented with 5% (v/v) glycerol. Monochromatic X-ray diffraction data were collected at 1.00 Å using beamline 22-ID of the Advanced Photon Source (Argonne National Laboratory) using an Eiger 16M detector. While several samples were harvested from this condition (and subsequent replications of this condition), only two samples diffracted X-rays beyond ∼10 Å and ultimately a single dataset was collected on one of these crystals. Diffraction data were integrated, scaled, and reduced using the HKL2000 software suite (*84*) and a high-resolution cut-off of 4.1 Å was chosen on the basis of I/σI and CC_1/2_ (**Table 1**). Consistent with the weak diffraction patterns observed, and difficulty in obtaining diffraction data in certain crystal orientations, PHENIX XTRIAGE (*85*) analysis suggested several data pathologies. This included severe anisotropy and low overall completeness. Additionally, strong evidence of translational non-crystallographic symmetry was observed by Patterson map analysis. Initial space group analysis suggested primitive tetragonal symmetry, however the final solution was obtained in space group *P* 21 21 2 (**Table S1**). Molecular replacement was carried out with PHASER in the Phenix software suite (*85*). Although translational non-crystallographic symmetry was detected, the option to account for this automatically during MR was deselected after initial failed solutions. An initial search was performed using a model of the monomeric activated form of recombinant C1r-CCP1-CCP2-SP (PDB: 2QY0, (*86*)) where four C-terminal loop residues were removed from Chain A. This search produced a solution in *P* 21 21 2 (LLG:248; TFZ: 18.9). This solution was fixed, and a monomeric copy of BBK32-C was used for a subsequent search producing a new solution (LLG: 424; TFZ: 17.8). Initial refinement of this solution using a single round of individual B-factors was performed (R_work_ = 42.5% / R_free_ = 44.4%) producing readily interpretable *2 F*o-*F*c density maps for both BBK32-C and C1r. Analysis of the asymmetric unit suggested two copies of the complex were probable (one copy: 74.3% solvent content / Matthews coeff.: 4.79 vs. two copies: 48.7% / 2.39). Additional exhaustive searches of various combinations of the complex, complex subcomponents, or individual domains yielded highly scoring solutions, however, presented unrealistic packing that could not be resolved by occupancy refinement. Higher and lower symmetry space groups were explored with no improvement. While we acknowledge other possibilities exist, our judgement is that two copies of the complex existed in the crystal, but the second copy cannot be confidently placed due to the underlying poor data quality and associated data pathologies of the crystal sample. Extensive efforts failed at producing better quality crystal samples, leading us to turn to an alternative SAXS-based approach. However, interpretable density of the single copy of the complex (**Fig. 1A, S1**) prompted us to carry forward with refinement in the absence of the second suspected asymmetric unit copy. Analysis of the contacts between BBK32-C and C1r in the obtained solution revealed three interfaces. Two of these interfaces were sized consistent with crystallographic contacts (346 Å^2^, and 344 Å^2^) and one large interface formed by a symmetry-mate of 1,108 Å^2^, which was rearranged into the asymmetric unit. Using this model, successive rounds of manual building using COOT (*81*) and iterative individual B-factor refinement using PHENIX.REFINE resulted in the final model 36.7%/37.2% (R_work_/ R_free_). A description of crystal cell constants, diffraction data quality, and properties of the final model can be found in **Table S1**. Representations of protein structures and electron density maps were generated in PyMol (www.pymol.org/).

### Surface Plasmon Resonance

SPR experiments were performed on a Biacore T200 instrument at 25° C at a flowrate of 30 µL min^-1^ with a running buffer of HBS-T (10 mM HEPES (pH 7.3), 140 mM NaCl, 0.005% Tween 20). Proteins were immobilized either on a NiD 200M Ni^2+^ chelator-modified sensorchips or CMD 200 biosensor chips (Xantec Bioanalytics) using standard amine coupling as previously described (*33*). For experiments involving immobilization of HIS-tagged species of C1r-CCP2-SP, NiD 200M surfaces were conditioned for 5 min with 0.5 M Na-EDTA (pH 8.5), followed by equilibration in HBS-T for 2 min. A solution of 5 mM nickel chloride was then injected for 2 min. Next, HIS-tagged ligands were captured on individual flow cells at densities between 300-400 RU. Single cycle injection curves were performed at five concentrations of BBK32-C or BBK32-C site-directed mutants (1, 3.2, 10, 32, 500 nM). For experiments using amine coupling, immobilization densities for each protein used in the study are presented in **Table S2**. Single cycle injection curves were performed with a five concentration five-fold injection series of (0-500 nM) C1r-CCP2-SP with 2 min association and a final dissociation time of 30 min. BBK32-C was immobilized under two immobilization densities on adjacent flow cells on the same chip: BBK32-C_High Density_ (2011.4 RU) and BBK32-C_Low Density_ (604.7 RU). Subsequent regeneration was performed by three 30 s injections of a regeneration solution (0.1 M glycine (pH 2.2), 2.5 M NaCl). Kinetic analysis was performed on each injection series using a 1:1 (Langmuir) binding model was applied using Biacore T200 Evaluation Software (GE Healthcare). Equilibrium dissociation constants (*K*_D_) were determined from resulting fits and are shown as mean *K*_D_. Quality of fits were assessed by inspection of residual plots (**Fig. S4. S5**).

### Classical pathway inhibitory assay

Classical pathway inhibition was determined using an ELISA-based assay as previously described (*33, 41, 87*). Briefly, high-binding 96 well plates (Grenier Bio-One) were coated with human IgM (MP Biomedical) and activation of classical pathway was monitored using 2% normal human serum (NHS; Innovative Research) and C4b detection was determined by monoclonal antibody (Santa Cruz Biotechnology). Dose-dependent inhibition was performed using a 12-point, two-fold dilution series of BBK32-C and mutants ranging from 1.1 to 2200 nM. Each assay was performed at least in duplicate and normalized to in-column positive (no inhibitor) and negative (no serum) controls.

### Serum complement sensitivity assay

*B. burgdorferi* strain B314 were grown at 32°C in 1% CO_2_ to early- to mid-log phase and diluted in complete BSK-II media to a concentration of 1×10^6^ cell/ml. *B. burgdorferi* cells were incubated in normal human serum (NHS; Complement Technology) at a final concentration of 15%. All strains were tested in duplicate independently four times. Samples were incubated at 37°C with gentle agitation for 1.5 hours with viability assessed by dark field microscopy.

Sensitivity was assessed by lack of motility, overt membrane damage, and/or cell lysis. The data is represented as the average of all fields counted for biological replicates (40 minimum) in each of the four independent assays.

### Immunoblots

Western immunoblots were conducted to detect BBK32 from the parent and mutant constructs produced in *B. burgdorferi* strain B314. Protein lysates were resolved by SDS-12.5% PAGE and transferred to PVDF membranes. Membranes were incubated with either a monoclonal antibody to BBK32 (diluted at 1:20,000) or *B. burgdorferi* strain B31 FlaB (Affinity BioReagents; diluted at 1:10,000), washed, and then incubated with a 1:10,000 dilution of Goat anti-mouse IgG-HRP (Thermo Fisher). The blots were processed to visualize the antigens as previously described (*41*)

### C1s proenzyme activation assay

Inhibition of C1r enzyme mediated activation of C1s proenzyme was assessed by a colorimetric peptide cleavage assay. 2.2 µM BBK32-C or R248A-K327A was incubated with 25 nM C1r enzyme, 100 µM DTNB (5,5’-Dithiobis-(2-Nitrobenzoic Acid), Ellman’s Reagent), and C1s specific active site analog 100 µM Z-L-Lys-thiobenzyl (MP Biomedical). Lastly 3.13 nM proenzyme C1s was added in HBS-T buffer (10 mM HEPES (pH7.3), 140 mM NaCl, .005% Tween 20). Absorbance was monitored at 405 nm at 37° C on a Versamax plate reader for 1 hr in triplicate. Data were in column normalized to no inhibitor and no C1s proenzyme controls.

### C1r peptide cleavage enzyme assays

C1r-CCP2-SP or C1r proenzyme by BBK32-C or R248A-K327A, a colorimetric peptide cleavage assay was employed (*55*). Single dose inhibition of 25 nM C1r-CCP2-SP or 25 nM C1r proenzyme with 2.2 µM BBK32-C or R248A-K327A was performed in the presence of 300 µM Z-G ly-Arg-thiobenzyl (MP Biomedical). Esterolytic cleavage was assayed by 100 µM DTNB (5,5’-Dithiobis-(2-Nitrobenzoic Acid), Ellman’s Reagent) in HBS-T buffer (10 mM HEPES (pH7.3), 140 mM NaCl, .005% Tween 20). Absorbance was monitored at 405 nm at 25° C on a Versamax plate reader for 1 hr in triplicate. Data were in column normalized to no inhibitor and no C1r-CCP2-SP controls, with 100% cleavage defined as the amount of Z-Gly-Arg-thiobenzyl cleaved by C1r-CCP2-SP in 1 hr at 25° C. Experimentation was performed in triplicate.

## Data Availability

SAXS data and models are deposited in the Small Angle Scattering Biological Data Bank (www.sasdb.org) with accession codes SASDKX7 (BBK32-C/C1r-CCP2-SP), SASDKV7 (C1r-CCP2-SP), and SASDKW7 (BBK32-C). The co-crystal structure of BBK32-C/C1r_300-705_ was deposited into the Protein Data Bank, Research Collaboratory for Structure Bioinformatics, Rutgers University (www.rcsb.org/) under the PDB code 7MZT.

## Acknowledgments

SAXS data was collected at SIBYLS which is supported by the DOE-BER IDAT DE-AC02-05CH11231 and NIGMS ALS-ENABLE (P30 GM124169 and S10OD018483). X-ray diffraction data were collected at Southeast Regional Collaborative Access Team 22-ID beamline at the Advanced Photon Source, Argonne National Laboratory. Supporting institutions may be found at www.ser-cat.org/members.html. Use of the Advanced Photon Source was supported by the US Department of Energy, Office of Science, Office of Basic Energy Sciences, under Contract W-31-109-Eng-38. We thank Dr. Brian Geisbrecht (Kansas State University) for helpful discussion related to this work. This study was supported by Public Health Service Grants AI133367 and AI146930 (to JTS and BLG).

