## Supplemental Materials for "A structural basis for inhibition of the complement initiator protease C1r by Lyme disease spirochetes"

Running Title: The molecular interface of BBK32/C1r

Ryan J. Garrigues[1], Alexandra D. Powell Pierce[2], Michal Hammel[3],
Jon T. Skare[2], *, and Brandon L. Garcia[1],*

[1]Department of Microbiology and Immunology, Brody School of Medicine, East Carolina University, Greenville, North Carolina, United States of America

[2]Department of Microbial Pathogenesis and Immunology, College of Medicine, Texas A&M University, Bryan/College Station, Texas, United States of America

[3]Molecular Biophysics and Integrated Bioimaging, Lawrence Berkeley National Laboratory, Berkeley, CA, USA

**This PDF file includes:**

Figs. S1 to S8
Tables S1 to S3

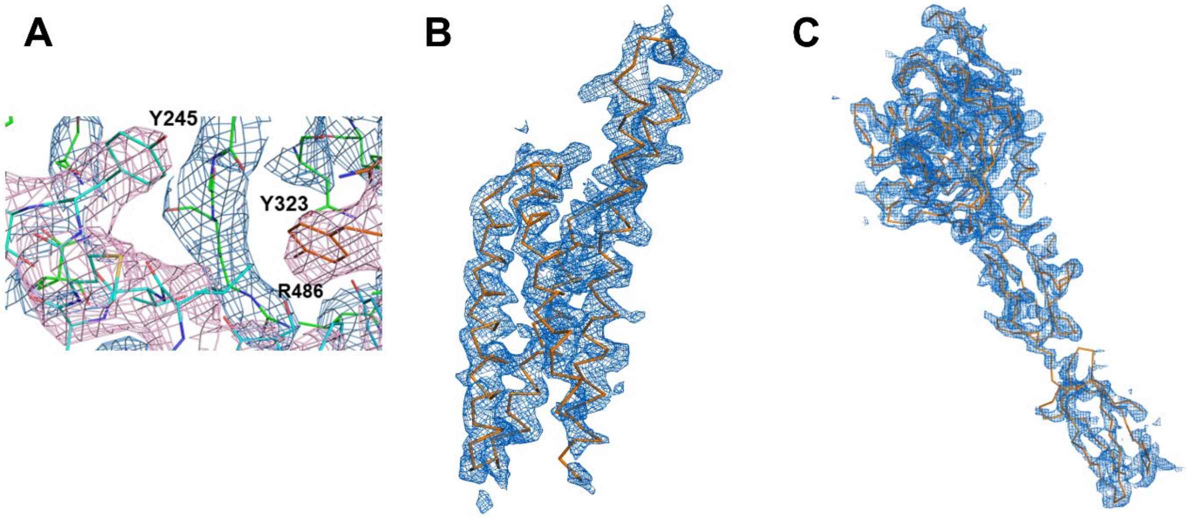

**Supplementary Figure S1. Electron density map quality.** 2*Fo-Fc* density contoured at 1.2 σ for **A)** Close-up view of representative area of the BBK32-C/C1r interface. **B)** BBK32-C (Chain: I) and **C)** C1r (Chains: A, B).

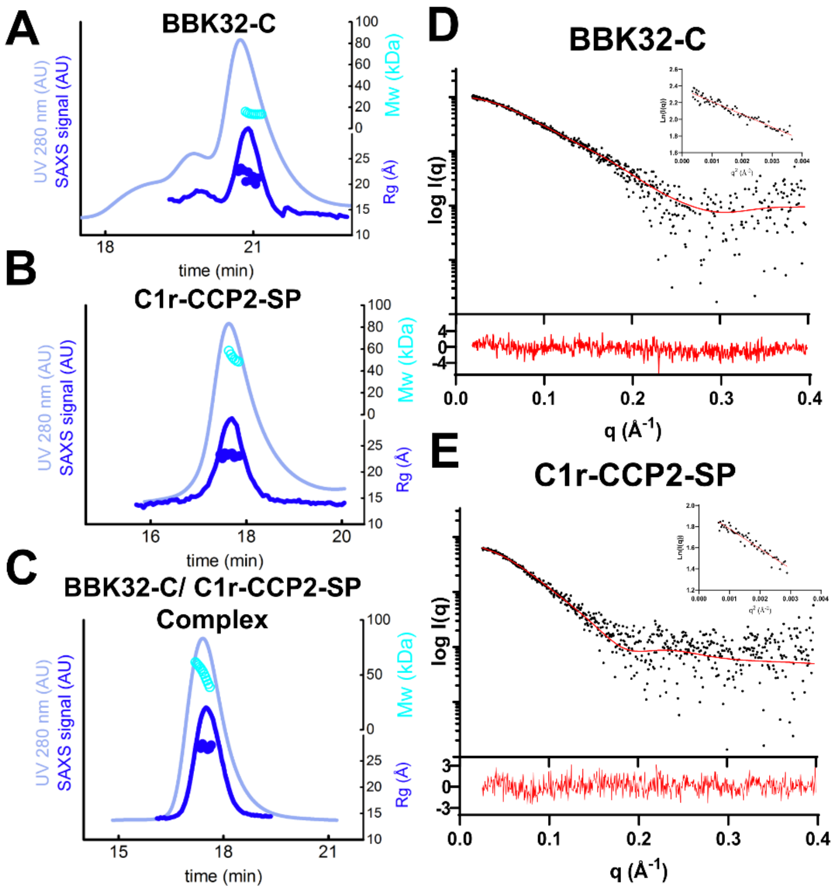

**Supplementary Figure S2. SEC-SAXS Data. A-C)** SEC-MALS-SAXS chromatographs for BBK32-C, C1r-CCP2-SP and BBK32-C/C1r-CCP2-SP. Solid lines represent the UV 280 nm (light blue) or integrated SAXS signal (dark blue) in arbitrary units, while symbols represent molecular mass (cyan) and Rg values for each collected SAXS frame (blue) versus elution time. **D-E)** Experimental SAXS curves for BBK32-C and C1r-CCP2-SP (black), respectively with FoXS calculated model curves (red). Residual error plots are presented below with Guinier plots (inset).

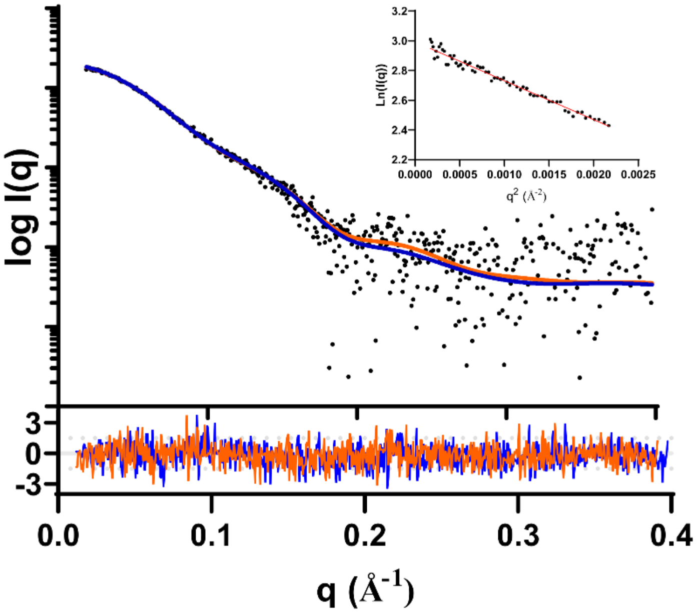

**Supplementary Figure S3. Comparison of the BBK32-C/C1r-CCP2-SP SAXS structure to the BBK32-C/C1r co-crystal structure.** Experimental SAXS curve for the C1r-CCP2-SP/BBK32-C complex sample (black) shown with calculated theoretical SAXS of the model (orange). The co-crystal of BBK32-C in complex with an autoproteolytic fragment of human C1r (PDB: 7MZT) where the CCP1 domain was removed was compared to the SAXS theoretical model curve obtained for the BBK32-C/C1r-CCP2-SP using FoXS. The calculated fit is shown in blue ($\chi^2$ = 1.20) with residual plot presented below for both fits. A Guinier plot is shown inset.

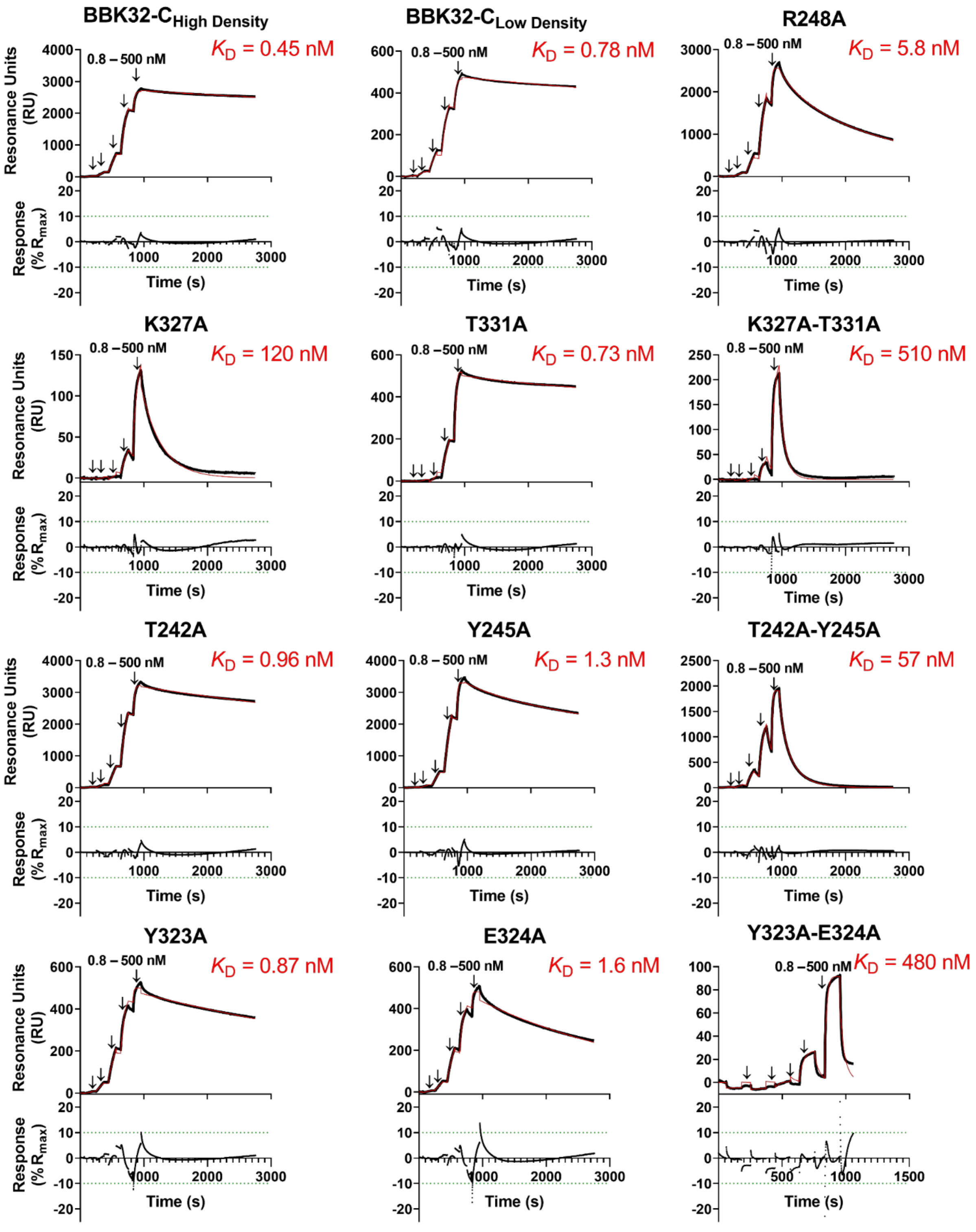

**Supplemental Figure S4. Kinetic fits of SPR binding assays with BBK32-C site-directed mutants**. Fitted sensorgrams of each curve presented in **Fig. 2C** with experimental data (black)

and a 1:1 Langmuir model fit (red). Downward arrows indicate injection phases of each C1r-CCP2-SP concentration in the series (0.8, 4.0, 20.0, 100, 500 nM). Residual plots for each SPR curve is normalized as a function of the fitted $R_{max}$ (**Table S2**). Dissociation constants ($K_D$) are shown as an average of duplicate experimentation. Additional SPR parameters for these experiments are shown in **Table S2.**

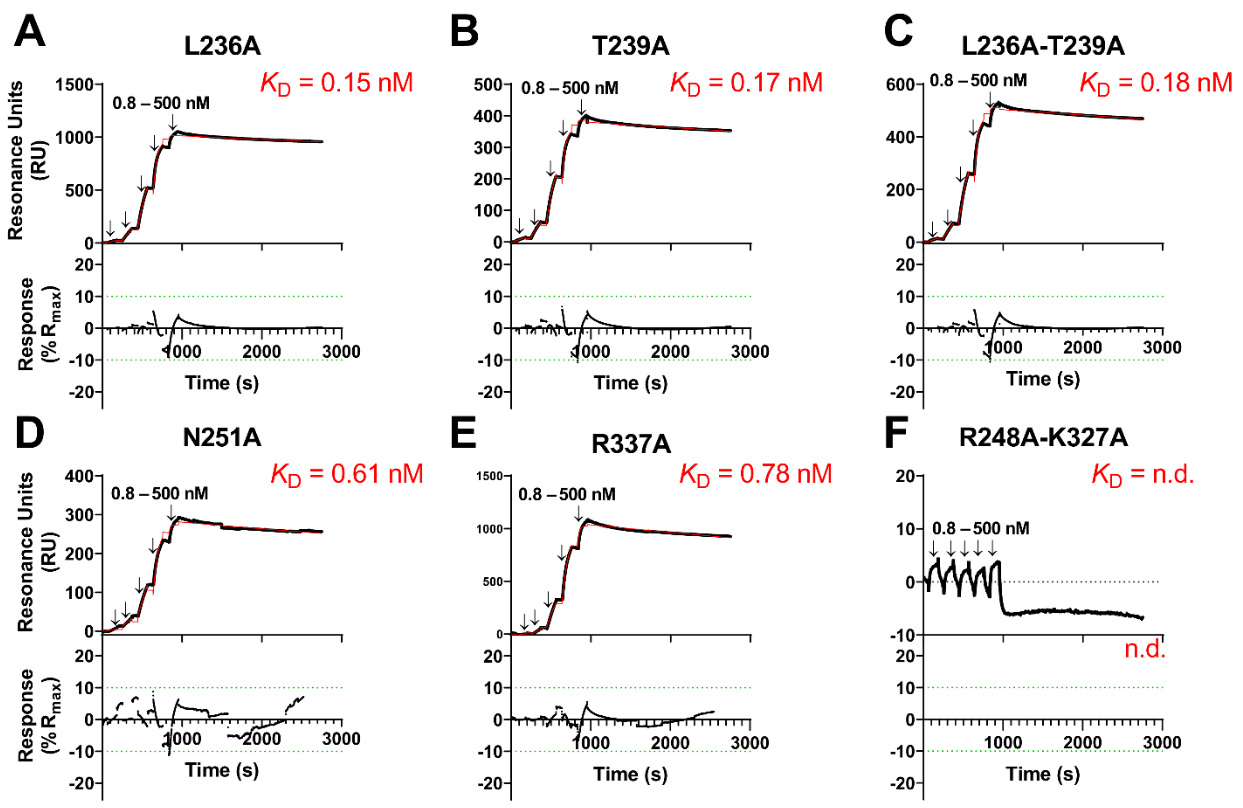

**Supplemental Figure S5. SPR binding assays with additional BBK32-C site-directed mutants**. **A-F)** A fivefold concentration series of C1r-CCP2-SP (0.8, 4.0, 20.0, 100, 500 nM) was injected over each immobilized BBK32-C mutant. Arrows indicate injection phases of each concentration. SPR sensorgrams (black) with a 1:1 Langmuir model fit (red) are shown. SPR experiments were performed in duplicate and dissociation constants ($K_D$) are shown. Residual plots are shown underneath each fitted curve and are normalized to calculated $R_{max}$ values (**Table S2**) **F)** Resulting curves were unable to be fit for BBK32-R248A-K327A and therefore do not include $K_D$ nor resulting residual plot. Additional SPR parameters for these curves are presented in **Table S2**.

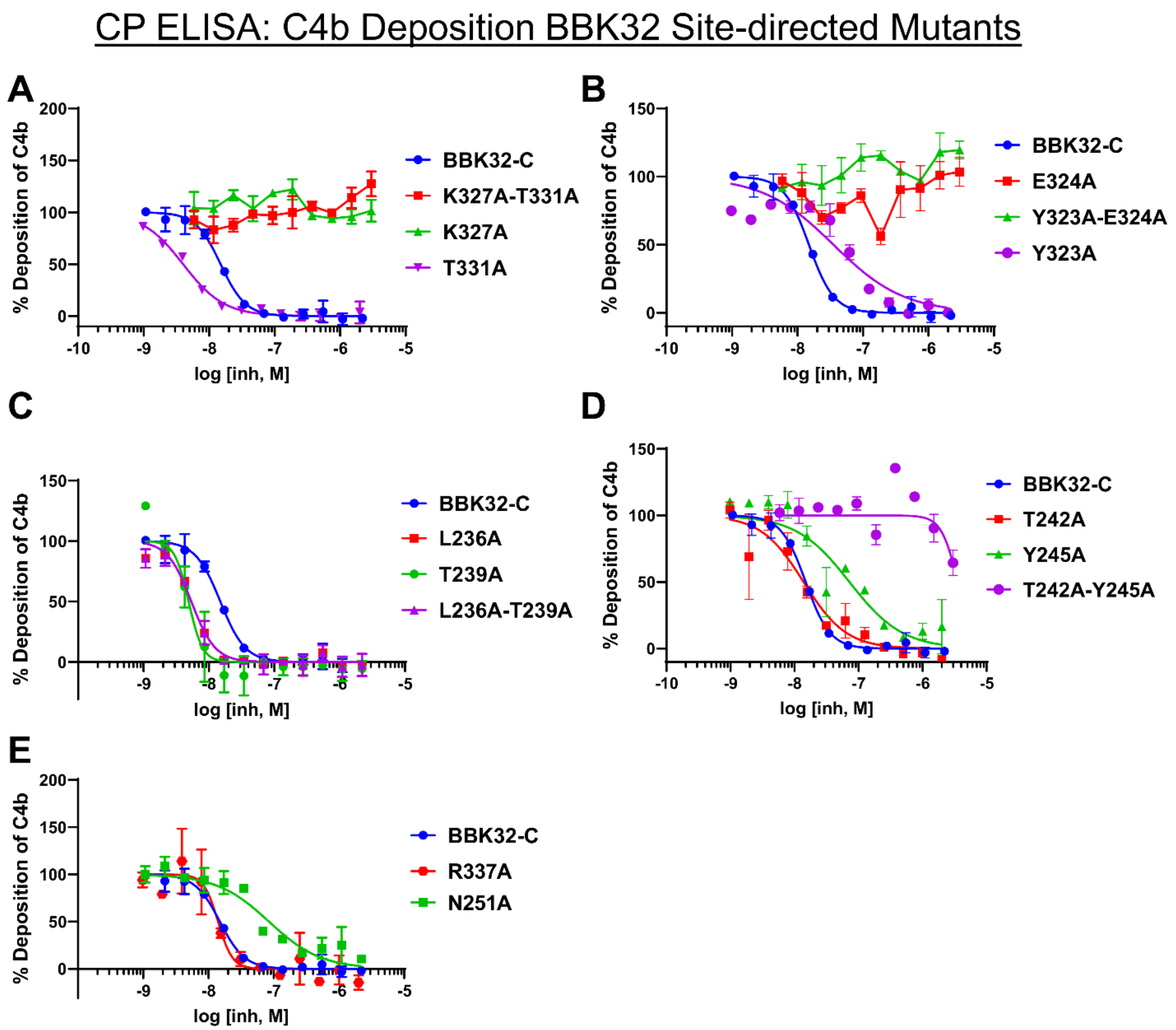

**Supplemental Figure S6. Complement inhibitory activity of additional BBK32 site-directed mutants.** Classical pathway ELISAs were performed to determine the ability of BBK32-C mutants to inhibit the classical pathway of complement in vitro. $IC_{50}$ and 95% confidence intervals are shown in **Table S1**. BBK32-C is shown on each plot for comparison only and is replotted from the BBK32-C data shown in **Fig. 4A**.

```
                    206                                       248
B. burgdorferi B31  SNRYQSYLEGVKYNVDSAIQTITKIYNTYTLFSTKLTQMYSTRLDNFAKAKAKEEAAKFTKEDLEKNFKTLL
WP_106021219.1      SNRYQSYLEGVKYNVDSAIQTITKIYNTYTLFSTKLTQMYSTRLDNFAKAKAKEEAAKFTKEDLEKNFKTLL
WP_010258321.1      SNRYQSYLEGVKYNVDSAIQTITKIYNTYTLFSTKLTQMYSTRLDNFAKAKAKEEAAKFTKEDLEKNFKTLL
WP_115655744.1      SNRYQSYLEGVKYNVDSAIQTITKIYNTYTLFSTKLTQMYSTRLDNFAKAKAKEEAAKFTKEDLEKNFKTLL
WP_012665899.1      SNRYQSYLEGVKYNVDSAIQTITKIYNTYTLFSTKLTQMYSTRLDNFAKAKAKEEAAKFTKEDLEKNFKTLL
WP_012666231.1      SNRYQSYLEGVKYNVDSAIQTITKIYNTYTLFSTKLTQMYSTRLDNFAKAKAKEEAAKFTKEDLEKNFKTLL
WP_106011580.1      SNRYQSYLEGVKYNVDSAIQTITKIYNTYTLFSTKLTQMYSTRLDNFAKAKAKEEAAKFTKEDLEKNFKTLL
WP_014540497.1      SNRYQSYLEGVKYNVDSAIQTITKIYNTYILFSTKLTQMYSTRLDNFAKAKAKEEAAKFTKEDLEKNFKTLL
WP_106022864.1      SNRYQSYLEGVKYNVDSAIQTITKIYNTYILFSTKLTQMYSTRLDNFAKAKAKEEAAKFTKEDLEKNFKTLL
WP_015056335.1      SNRYQSYLEGVKYNVDSAIQTITKIYNTYILFSTKLTQMYSTRLDNFAKAKAKEEAAKFTKEDLEKNFKTLL
WP_012622643.1      SNRYQSYLEGVKYNVDSAIQTITKIYNTYILFSTKLTQMYSTRLDNFAKAKAKEEAAKFTKEDLEKNFKTLL
WP_106058032.1      SNRYQSYLEGVKYNVDSAIQTITKIYNTYTLFSTKLTQMYSTRLDNFAKAKAKEEAAKFTKEDLEKNFKTLL
WP_015899365.1      SNRYQSYLEGVKYNVDSAIQTITKIYNTYILFSTKLTQMYSTRLDNFAKAKAKEEAAKFTKEDLEKNFKTLL
WP_106022719.1      SNRYQSYLEGVKYNVDSAIQTITKIYNTYTLFSTKLTQMYSTRLDNLTKAKAKEEAAKFTKEDLEKNFKTLL
WP_106017555.1      SNRYQSYLEGVKYNVDSAIQTITKIYNTYTLFSTKLTQMYSTRLDNLTKAKAKEEAAKFTKEDLEKNFKTLL
WP_012614929.1      SNRYQSYLEGVKYNVDSAIQTITKIYNTYILFSTKLTQMYSTRLDNFAKAKAKEEAAKFTKEDLEKNFKTLL
AAL84596.1          SNRYQSYLEGVKYNVDSAINTINKIYDTYTLFSTKLTQMYSTRLDNLAKAKAKEEAAKFTKEDLEKNFKTLL
AAB71755.1          SNRYQSYLEGVKYNVDSAIQTITKIYNTYTLFSTKLTQMYSTRLDNFAKAKAKEEAAKFTKEDLEKNFKTLL
WP_106017066.1      SNRYQSYLEGVKYNVDSAIQTITKIYNTYTLFSTKLTQMYSTRLDNFAKAKAKEEAAKFTKEDLEKNFKTLL
WP_106023546.1      SNRYQSYLEGVKYNVDSAIQTITKIYNTYTLFSTKLTQMYSTRLDNFAKAKAKEEAAKFTKEDLEKNFKTLL
WP_014540529.1      SNRYQSYLEGVKYNVDSAIQTITKIYNTYTLFSTKLTQMYSTRLDNFAKAKAKEEAAKFTKEDLEKNFKTLL
WP_096890875.1      SNRYQSYLEGVKYNVDSAIQTITKIYNTYTLFSTKLTQMYSTRLDNFTKAKAKEEAAKFTKEDLEKNFNTLL
WP_012665967.1      SNRYQSYLEGVKYNVDSAIQTITKIYNTYTLFSTKLTQMYSTRLDNFAKAKAKEEAAKFTKEDLEKNFNTLL
WP_106017344.1      SNRYQSYLEGVKYNVDSAIQTITKIYNTYTLFSTKLTQMYSTRLDNFAKAKAKEEAAKFTKEDLEKNFNTLL
WP_106034380.1      SNRYQSYLEGVKYNVDSAIQTITKIYNTYTLFSTKLTQMYSTRLDNFAKAKAKEEAAKFTKEDLEKNFNTLL
                    *******************:**.***:** ****************::********************:***

                                                                  324 327              348
B. burgdorferi B31  NYIQVSVKTAANFVYINDTHAKRKLENIEAEIKTLIAKIKEQSNLYEAYKAIVTSILLMRDSLKEVQGIID
WP_106021219.1      NYIQVSVKTAANFVYINDTHAKRKLENIEAEIKTLIAKIKEQSNLYEAYKAIVTSILLMRDSLKEVQGIID
WP_010258321.1      NYIQVSVKTAANFVYINDTHAKRKLENIEAEIKTLIAKIKEQSNLYEAYKAIVTSILLMRDSLKEVQGIID
WP_115655744.1      NYIQVSVKTAANFVYINDTHAKRKLENIEAEIKTLIAKIKEQSNLYEAYKAIVTSILLMRDSLKEVQGIID
WP_012665899.1      NYIQVSVKTAANFVYINDTHAKRKLENIEAEIKTLIAKIKEQSNLYEAYKAIVTSILLMRDSLKEVQGIID
WP_012666231.1      NYIQVSVKTAANFVYINDTHAKRKLENIEAEIKTLIAKIKEQSNLYEAYKAIVTSILLMRDSLKEVQGIID
WP_106011580.1      NYIQVSVKTAANFVYINDTHAKRKLENIEAEIKTLIVKIKEQSNLYEAYKAIVTSILLMRDSLKEVQGIID
WP_014540497.1      NYIQVSVKTAANFVYINDTHAKRKLENIEAEIKTLIAKIKEQSNLYEAYKAIVTSILLMRDSLKEVQGIID
WP_106022864.1      NYIQVSVKTAANFVYINDTHAKRKLENIEAEIKTLIAKIKEQSNLYEAYKAIVTSILLMRDSLKEVQGIID
WP_015056335.1      NYIQVSVKTAANFVYINDTHAKRKLENIEAEIKTLIAKIKEQSNLYEAYKAIVTSILLMRDSLKEVQGIID
WP_012622643.1      NYIQVSVKTAANFVYINDTHAKRKLENIEAEIKTLIAKIKEQSNLYEAYKAIVTSILLMRDSLKEVQGIID
WP_106058032.1      NYIQVSVKTAANFVYINDTHAKRKLENIEAEIKTLITKIKEQSNLYGAYKAIVTSILLMRDSLKEVQGIID
WP_015899365.1      NYIQVSVKTAANFVYINDTHAKRKLENIEAEIKTLIAKIKEQSNLYGAYKAIVTSILLMRDSLKEVQGIID
WP_106022719.1      NYIQVSVKTAANFVYINDTHAKRKLENIEAEIKTLIVKIKEQSNLYEAYKAIVTSILLMRDSLKEVQGIID
WP_106017555.1      NYIQVSVKTAANFVYINDTHAKRKLENIEAEIKTLITKIKEQSNLYEAYKAIVTSILLMRDSLKEVQGIID
WP_012614929.1      NYIQVSVKTAANFVYINDTHAKRKLENIEAEIKTLITKIKEQSNLYGAYKAIVTSILLMRDSLKEVQGIID
AAL84596.1          NYIQVSVKTAANFVYINDTHAKRKLENIEAEIKTLIAKIKEQSNLYEAYKAIVTSILLMRDSLKEVQGIID
AAB71755.1          NYIQVSVKTAANFVYINDTHAKRKLENIETEIKTLIAKIKEKPDLYQAYKAIVTPILLMRDSLKEVQSAID
WP_106017066.1      NYIQVSVKTAANFVYINDTHAKRKLENIETEIKTLIAKIKEKPDLYQAYKAIVTPILLMRDSLKEVQSAID
WP_106023546.1      NYIQVSVKTAANFVYINDTHAKRKLENIETEIKTLIAKIKEKPDLYQAYKAIVTPILLMRDSLKEVQSAID
WP_014540529.1      NYIQVSVKTAANFVYINDTHAKRKLENIETEIKTLIAKIKEKPDLYQAYKAIVTPILLMRDSLKEVQSAID
WP_096890875.1      NYIQVSVKTAANFVYINDTHAKRKLENIETEIKTLIAKIKEKPDLYQAYKAIVTPILLMRDSLKEVQSAID
WP_012665967.1      NYIQVSVKTAANFVYINDTHAKRKLENIETEIKTLIAKIKEKPDLYQAYKAIVTPILLMRDSLKEVQSAID
WP_106017344.1      NYIQVSVKTAANFVYINDTHAKRKLENIETEIKTLIAKIKEKPDLYQAYKAIVTPILLMRDSLKEVQSAID
WP_106034380.1      NYIQVSVKTAANFVYINDTHAKRKLENIETEIKTLIAKIKEKPDLYQAYKAIVTPILLMRDSLKEVQSAID
                    *****************************:******.****: :** ******* ************. **
```

**Supplemental Figure S7. Alignment of the BBK32-C region of *B. burgdorferi* BBK32 sequences.** NCBI BLASTp was used to query all *B. burgdorferi* genomes (taxid: 139) for BBK32 using the B31 strain sequence. The top 25 non-redundant sequences were aligned using Clustal

Omega (EMBL-EMI). Numbering is based on *B. burgdorferi* B31 BBK32-C, and residues R248, E324, and K327 are highlighted. R248 and K327 are conserved amongst all 25 sequences whereas E324 is substituted for a Gln or a Gly in 44% of the sequences shown.

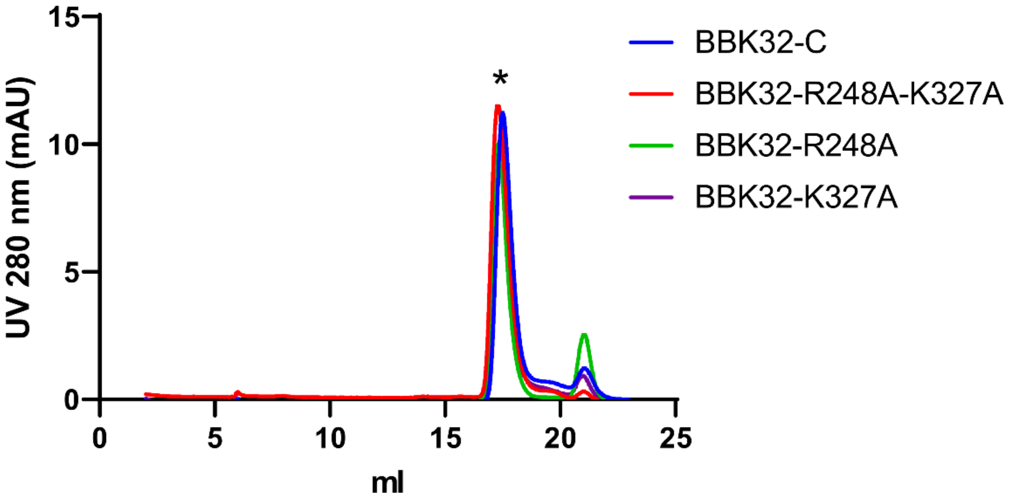

**Supplemental Figure S8. Gel filtration analysis of all mutants used in the study.** Gel filtration analysis on a Superdex 200 Increase 10/300 GL shows monodisperse peaks of similar elution volumes, indicating similar tertiary structure of BBK32-C mutants compared to wild type. Single peaks for BBK32-C and site-directed mutants between 17.3 and 17.5 mLs were observed (denoted with *).

**Table S1. X-ray Data collection and refinement statistics (molecular replacement)**

| | $C1r_{(300\text{-}705)}/BBK32_{(206\text{-}348)}$ |
|---|---|
| **Data collection** | |
| **Space group** | P 21 21 2 |
| **Cell dimensions** | |
| ***a, b, c,* Å** | 113.62, 96.99, 108.33 |
| **α, β, γ, °** | 90.00, 90.00, 90.00 |
| **Resolution, Å** | 50.0 – 4.10 (4.25 – 4.10) |
| $R_{pim}$ | 0.164 (0.264) |
| $CC_{1/2}$ | 0.997 (0.903) |
| $I/\sigma I$ | 4.4 (2.2) |
| ***No. of reflections*** | 15,139 (1,455) |
| **Completeness, %** | 85.5 (83.4) |
| **Redundancy** | 4.9 (4.3) |
| **Refinement** | |
| $R_{work}/R_{free}$ **(%)** | 36.9/37.2 |
| **No. non-hydrogen atoms** | 4,120 |
| ***B*-factors (range and mean)** | 59.13-185.56 (108.30) |
| **Root mean square deviations** | |
| **Bond lengths, Å** | 0.008 |
| **Bond angles, °** | 1.29 |

### Table S2. SPR and ELISA parameters

| | | Surface Plasmon Resonance | | | | CP ELISA | |
|---|---|---|---|---|---|---|---|
| **Ligand** | **Immobilization Density (RU)** | ***k*a (1/Ms)** | ***k*d (1/s)** | ***K*D (nM)** | **Rmax (RU)** | **IC50 (nM)** | **95% Confidence Interval (nM)** |
| **BBK32-$C_{High\ Density}$** | 2011.4 | 1.0 x $10^{5}$ | 4.5 x $10^{-5}$ | 0.45 ± 0.00 | 2717.2, 2680.2 | 15 | 13 to 17 |
| **BBK32-$C_{Low\ Density}$** | 604.7 | 8.1 x $10^{4}$ | 6.3 x $10^{-5}$ | 0.78 ± 0.03 | 476.3, 467.6 | | |
| **E324A** | 1098.8 | 2.0 x $10^{5}$ | 3.4 x $10^{-4}$ | 1.6 ± 0.07 | 441.3, 430.3 | n.d. | n.d. |
| **K327A** | 1087.5 | 9.7 x $10^{4}$ | 7.5 x $10^{-3}$ | 120 ± 50 | 185.6, 149.3 | n.d. | n.d. |
| **K327A-T331A** | 938.4 | 3.7 x $10^{4}$ | 2.6 x $10^{-2}$ | 510 ± 200 | 479.8, 321.6 | n.d. | n.d. |
| **L236A** | 764.6 | 2.3 x $10^{5}$ | 3.5 x $10^{-5}$ | 0.15 ± 0.00 | 1014, 1006 | 5.4 | 4.7 to 6.2 |
| **L236A-239A** | 305.0 | 2.4 x $10^{5}$ | 4.3 x $10^{-5}$ | 0.18 ± 0.00 | 504.4, 500.0 | 5.4 | 4.7 to 6.2 |
| **N251A** | 1169.9 | 1.8 x $10^{5}$ | 1.1 x $10^{-4}$ | 0.61 ± 0.21 | 282.8, 281.7 | 85 | 59 to 120 |
| **R248A** | 2564.4 | 2.2 x $10^{5}$ | 1.3 x $10^{-3}$ | 5.8 ± 0.10 | 2576, 2559 | 420 | 280 to 630 |
| **R248A-327A** | 1222.1 | n.d. | n.d. | n.d. | n.d. | n.d. | n.d. |
| **R337A** | 1538.8 | 1.1 x $10^{5}$ | 8.8 x $10^{-5}$ | 0.78 ± 0.13 | 1070, 1041 | 14 | 11 to 18 |
| **T239A** | 250.7 | 2.6 x $10^{5}$ | 4.4 x $10^{-5}$ | 0.17 ± 0.00 | 379.8, 377.1 | 4.9 | 4.0 to 6.2 |
| **T242A** | 2837.4 | 1.0 x $10^{5}$ | 9.9 x $10^{-5}$ | 0.96 ± 0.00 | 3192, 3157 | 14 | 9.2 to 20.9 |
| **T242A-245A** | 2055.4 | 2.1 x $10^{5}$ | 1.2 x $10^{-2}$ | 57 ± 0.10 | 2117, 2093 | 3600 | 2700 to 12000 |
| **T331A** | 940.3 | 1.4 x $10^{5}$ | 8.2 x $10^{-5}$ | 0.73 ± 0.30 | 502.6, 516.5 | 4 | 3.6 to 4.5 |
| **Y245A** | 2706.6 | 2.8 x $10^{5}$ | 3.5 x $10^{-4}$ | 1.3 ± 0.01 | 3334, 3310 | 72 | 44 to 120 |

| | | | | | | | |
|---|---|---|---|---|---|---|---|
| **Y323A** | 1058.1 | 1.8 x $10^5$ | 1.6 x $10^{-4}$ | 0.87 ± 0.06 | 475.1, 461.0 | 36 | 21 to 56 |
| **Y323A-324A** | 1496.9 | 1.0 x $10^5$ | 4.6 x $10^{-2}$ | 480 ± 51 | 146.5, 120.5 | n.d. | n.d. |

n.d. not determined

**Table S3. Strains, plasmid constructs, and oligonucleotides used in this study.**

| | | |
|---|---|---|
| ***E. coli*** **strains** | | |
| NEB® 5-alpha | *fhuA2* (*argF-lacZ*)U169 *phoA glnV44* 80(*lacZ*)ΔM15 *gyrA96 recA1 relA1 endA1 thi-1 hsdR17* | New England Biolabs |
| ***B. burgdorferi*** **strains** | | |
| B314 | Serum-sensitive, non-infectious *B. burgdorferi* B31 derivative strain lacking all linear plasmids | *(56)* |
| B314 pBBE22*luc* | Serum-sensitive, non-infectious *B. burgdorferi* B31 derivative strain lacking all linear plasmids; shuttle vector encodes *bbe22* and *B. burgdorferi* codon optimized *luc* gene under the control of a strong borrelial promoter ($P_{flaB}$-*luc*). | *(74)* |
| B314 pCD100 | Serum-sensitive, non-infectious *B. burgdorferi* B31 derivative strain lacking all linear plasmids with wildtype *bbk32* under control of its native promoter in pBBE22*luc*. | This study |
| B314 pAP5 | Serum-sensitive, non-infectious *B. burgdorferi* B31 derivative strain lacking all linear plasmids with *bbk32* R248A under control of the *bbk32* promoter in pBBE22*luc.* | This study |
| B314 pAP6 | Serum-sensitive, non-infectious *B. burgdorferi* B31 derivative strain lacking all linear plasmids with *bbk32* K327A under control of the *bbk32* promoter in pBBE22*luc*. | This study |
| B314 pAP7 | Serum-sensitive, non-infectious *B. burgdorferi* B31 derivative strain lacking all linear plasmids with *bbk32* R248A/K327A under control of the *bbk32* promoter in pBBE22*luc*. | This study |
| **Plasmids** | | |
| pBBE22*luc* | Borrelial shuttle vector containing *bbe22* and *B. burgdorferi* codon-optimized *luc* gene under the control of a strong borrelial promoter ($P_{flaB}$-*luc*) | *(74)* |
| pCD100 | Knock in construct of wild-type BBK32 with its native promoter in pBBE22*luc*. | *(33)* |

| | | |
|---|---|---|
| pAP5 | Knock in construct encoding BBK32 R248A under the control of its native *bbk32* promoter in pBBE22*luc*. | This study |
| pAP6 | Knock in construct encoding BBK32 K327A under the control of its native BBK32 promoter in pBBE22*luc*. | This study |
| pAP7 | Knock in construct encoding BBK32 R248A-K327A under the control of its native BBK32 promoter in pBBE22*luc*. | This study |

| **Oligonucleotides** | | | |
|---|---|---|---|
| R248A mutant F | GTATTCTACAGCACTTGACAACTTTGCTAAAGCC | Oligonucleotide pair used for site-directed mutagenesis of *bbk32* R248 for conversion to alanine and generate pAP5 | This study |
| R248A mutant R | AGTTGTCAAGTGCTGTAGAATACATTTGGGTTAGCTTTGT | | |
| *pncAF* | TATTGGAATTAATAGGCGGTGATG | Oligonucleotide pair used to confirm *bbk32* knock-in constructs | *(41)* |
| *lucF* | GAGGGGTTGTATTTGTTGACG | | |
| K327A frag F | AAGTAAGTGTAAAGACTGCAGCAAATTTTGTATACATA | Oligonucleotide pair used to amplify the *bbk32* K327A fragment for recombination with either pCD100 (to make pAP6) or pAP5 (to make pAP7). | This study |
| K327A frag R | TATGTATACAAAATTTGCTGCAGTCTTTACACTTACTT | | |
| backbone frag F | GGCTACATAATATGTCGACCTGCAGGCATGCAAGCTT | Oligonucleotide pair used to | This study |

| | | |
|---|---|---|
| backbone frag R | AAGCTTGCATGCCTGCAGGTCGAC ATATTATGTAGCC | amplify either pCD100 (to make pAP6) or pAP5 (to make pAP7) fragments for recombination with the K327A fragment. |
